# Dynamics of genome architecture and chromatin function during human B cell differentiation and neoplastic transformation

**DOI:** 10.1101/764910

**Authors:** Roser Vilarrasa-Blasi, Paula Soler-Vila, Núria Verdaguer-Dot, Núria Russiñol, Marco Di Stefano, Vicente Chapaprieta, Guillem Clot, Irene Farabella, Pol Cuscó, Xabier Agirre, Felipe Prosper, Renée Beekman, Silvia Beà, Dolors Colomer, Henk Stunnenberg, Ivo Gut, Elias Campo, Marc A. Marti-Renom, José Ignacio Martin-Subero

**Author notes:** These authors have contributed equally (Roser Vilarrasa-Blasi, Paula Soler-Vila). These authors have jointly supervised this work (Marc A. Marti-Renom, José Ignacio Martin-Subero).

## Abstract

Despite recent advances, the dynamics of genome architecture and chromatin function during human cell differentiation and its potential reorganization upon neoplastic transformation remains poorly characterized. Here, we integrate *in situ* Hi-C and nine additional omic layers to define and biologically characterize the dynamic changes in three-dimensional (3D) genome architecture across normal B cell differentiation and in neoplastic cells from different subtypes of chronic lymphocytic leukemia (CLL) and mantle cell lymphoma (MCL) patients. Beyond conventional active (A) and inactive (B) compartments, an integrative analysis of Hi-C data reveals the presence of a highly-dynamic intermediate compartment enriched in poised and polycomb-repressed chromatin. During B cell development, we detect that 28% of the compartments change at defined maturation stages and mostly involve the intermediate compartment. The transition from naive to germinal center B cells is associated with widespread chromatin activation, which mostly reverts into the naive state upon further maturation of germinal center cells into memory B cells. The analysis of CLL and MCL neoplastic cells points both to entity and subtype-specific alterations in chromosome organization. Remarkably, we observe that large chromatin blocks containing key disease-specific genes alter their 3D genome organization. These include the inactivation of a 2Mb region containing the *EBF1* gene in CLL and the activation of a 6.1Mb region containing the *SOX11* gene in clinically aggressive MCL. This study indicates that 3D genome interactions are extensively modulated during normal B cell differentiation and that the genome of B cell neoplasias acquires a tumor-specific 3D genome architecture.

## Introduction

Over the last decades, our understanding of higher-order chromosome organization in the eukaryotic interphase nucleus and its regulation of cell state, function, specification and fate has profoundly increased^1, 2^.

Chromatin conformation capture techniques have been used to elucidate the genome compartmentalization^3, 4^. It is widely accepted that the genome is segregated into two large compartments, named A-type and B-type^5^, which undergo widespread remodeling during cell differentiation^2, 6–9^. These compartments have been associated with different GC content, DNAsel hypersensitivity, gene density, gene expression, replication time, and chromatin marks^5, 10^. Alternative subdivisions of genome compartmentalization have been proposed, including three compartments^11^ or even five compartment subtypes with distinct genomic and epigenomic features^12^. All of these studies highlight the role of genome three-dimensional (3D) organization in the regulatory decisions associated with cell fate. However, the majority of these studies have been performed using cell lines, animal models or cultured human cells^7, 8, 13–15^, and although few analyze sorted cells from healthy human individuals^16, 17^, there is limited information regarding 3D genome dynamics across the differentiation program of a single human cell lineage^16^.

Normal human B cell differentiation is an ideal model to study the dynamic 3D chromatin conformation during cell maturation, as these cells show different transcriptional features and biological behaviors, and can be accurately isolated due to their distinct surface phenotypes^18, 19^. Moreover, how the 3D genome is linked to cancer development using primary samples from patients is also widely unknown^20^. In this context, several types of neoplasms can originate from B cells at distinct differentiation stages^21^. Out of them, chronic lymphocytic leukemia (CLL) and mantle cell lymphoma (MCL) are derived from mature B cells and show a broad spectrum of partially overlapping biological features and clinical behaviors^22^. Both diseases can be categorized according to the mutational status of the immunoglobulin heavy chain variable region (IGHV), a feature that seems to be related to the maturation stage of the cellular origin^23^. CLL cases lacking IGHV somatic hypermutation are derived from germinal center-independent B cells whereas CLL with mutated IGHV derive from germinal center-experienced B cells^24^. In CLL, this variable is strongly associated with the clinical features of the patients, with mutated IGHV (mCLL) cases correlating with good prognosis and those lacking IGHV mutation (uCLL) with poorer clinical outcome^24^. In MCL, although two groups based on the IGHV mutational status can be recognized and partially correlate with clinical behavior, other markers such as expression of the *SOX11* oncogene are used to classify cases into clinically-aggressive conventional MCL (cMCL) and clinically-indolent non-nodal leukemic MCL (nnMCL)^22, 25–27^.

From an epigenomic perspective, previous reports have identified that B cell maturation and neoplastic transformation to CLL or MCL entails extensive modulation of the DNA methylome and histone modifications^28–33^. However, whether such epigenetic changes are also linked to modulation of the higher-order chromosome organization is yet unknown^34^.

Here, to decipher the 3D genome architecture of normal and neoplastic B cells, we generated *in situ* high-throughput chromosome conformation capture (Hi-C) maps of cell subpopulations spanning the B cell maturation program as well as of neoplastic cells from MCL and CLL patients. Next, we mined the data together with whole-genome maps of six different histone modifications, chromatin accessibility, DNA methylation, and gene expression obtained from the same human cell subpopulations and patient samples. This multi-omics approach allowed us to identify a widespread modulation of the chromosome organization during human B cell maturation and neoplastic transformation, including the presence of recurrent aberrations in the chromosome organization of regions containing deregulated disease-specific genes.

## Results

### Multi-omics analysis during human B cell differentiation

We used *in situ* Hi-C to generate genome-wide chromosome conformation maps of normal human B cells across their maturation program. These included three biological replicates each of naive B cells (NBC), germinal center B cells (GCBC), memory B cells (MBC), and plasma cells (PC) (Figure 1A-1B and **Supplementary Table** 1). From the same B cell subpopulations, we analyzed nine additional omics layers generated as part of the BLUEPRINT consortium^28, 35^. Specifically, we obtained data for chromatin immunoprecipitation with massively parallel sequencing (ChIP-seq) of six histone modifications with non-overlapping functions (H3K4me3, H3K4me1, H3K27ac, H3K36me3, H3K9me3, H3K27me3), transposase-accessible chromatin with high-throughput sequencing (ATAC-seq), whole genome bisulfite sequencing (WGBS), and gene expression (RNA-seq).

**Figure 1.**
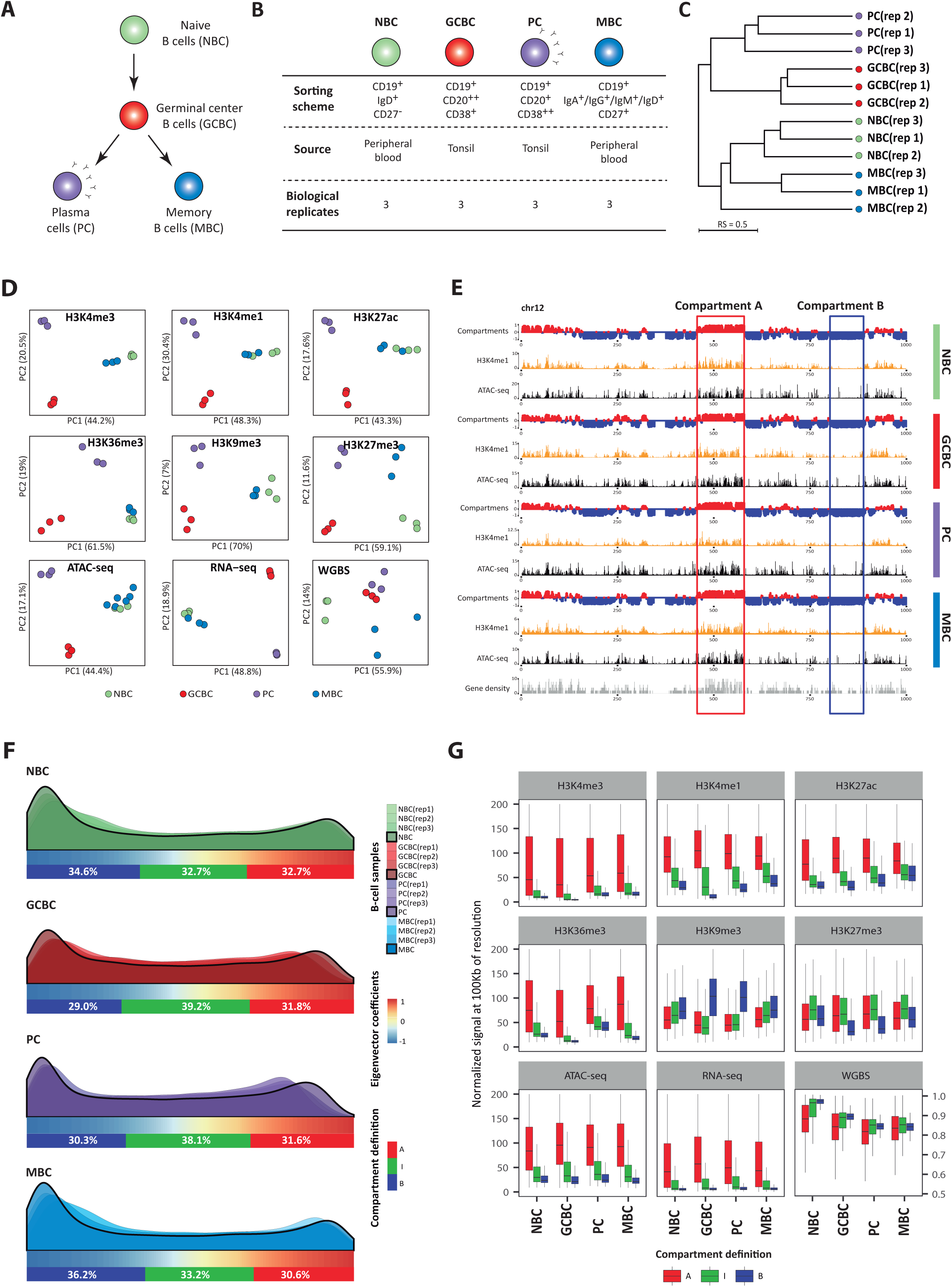
Multi-omics view of B cell differentiation and identification of an intermediate compartment. **A** - Schematic overview of mature B cell differentiation showing the four B cell subpopulations considered in this study. **B -** Sample description and *in situ* Hi-C sequencing experimental design for normal B cell differentiation subpopulations. NBC, naive B cells; GCBC, germinal center B cells; MBC, memory B cells and PC, plasma cells. **C** - Dendrogram of the reproducibility score of B cell subpopulation replicates for normalized Hi-C contact maps at 100Kb resolution. **D** - Unsupervised principal component analysis (PCA) for nine omics layers: chromatin immunoprecipitation followed by sequencing (ChIP-seq) of six histone marks (H3K4me3 n=46,184 genomic regions, H3K4me1 n=44,201 genomic regions, H3K27ac n=72,222 genomic regions, H3K36me3 n=25,945 genomic regions, H3K9me3 n=40,704 genomic regions, and H3K27me3 n=20,994 genomic regions), chromatin accessibility measured by ATAC-seq (n=99,327 genomic regions), DNA methylation measured by whole-genome bisulfite sequencing (WGBS, n=15,089,887 CpGs) and gene expression measured by RNA-seq (n=57,376 transcripts). Three independent biological replicates of NBC, GCBC, PC, and MBC were studied for all omic layers, with the exception of ATAC-seq for which six biological replicates of MBC were used. **E** - Example on chromosome 12 (chrl2) comparing the profile of three-dimensional (3D) data *(in situ* Hi-C), H3K4me1 ChIP-seq signal, chromatin accessibility (ATAC-seq) and gene density. The red and blue rectangles highlight the features of A and B compartments, respectively. **F -** Distribution of the first eigenvector of each B cell subpopulation (three replicates and merge). The relative abundance of A-type, B-type and intermediate (l)-type compartments per merged B cell subpopulations are indicated below each distribution. Compartment definition based on eigenvalue thresholds: A-type, 1 to 0.43; I-type, 0.43 to −0.63; B-type, −0.63 to −1. **G -** Boxplots showing the association between the three compartments (A-type, I-type and B-type) and each of the nine additional omics layers under study.

We initially explored the intra- and inter-subpopulation variability and observed that the Hi-C replicas were concordant, as quantified measuring and clustering the reproducibility score (RS)^36^ (Figure 1C and **Extended Data Figure 1A).** Furthermore, the comparison of samples suggests that the overall genome architecture of NBC is more similar to MBC, and clearly different from GCBC and PC, which belong to a different cluster (Figure 1C). This finding was also reflected in the first component of the principal component analysis (PCA) of histone modifications, chromatin accessibility and gene expression (Figure 1D). In contrast to other omics marks, the first component of DNA methylation data resulted in a division of GCBC, MBC and PC separated from the NBC. These analyses suggest fundamental differences between chromatin-based epigenetic marks, including chromosome conformation data, and DNA methylation. In fact, changes in DNA methylation linearly accumulate throughout B cell maturation^30,31^, which explains the clear differences between NBC and MBC in spite of their converging transcriptomes.

### Polycomb-associated chromatin defines an intermediate and moldable 3D genome compartment

To study the compartmentalization of the genome during B cell differentiation, we next merged all biological replicates per B cell subpopulation resulting in interaction Hi-C maps with around 300 million valid reads each. These Hi-C interaction maps were further segmented into positive and negative eigenvalues based on the eigenvector decomposition^5, 37^, and regions were assigned to the A-type (active) and B-type (inactive) compartments using the association with histone modifications (Figure 1E and **Extended Data Figure 1B).** A pairwise correlation of the first eigenvector of each B cell subpopulation showed that NBC and MBC on the one hand, and GCBC and PC on the other hand, have similar compartmentalization **(Extended Data Figure 1C**), confirming previous results using the RS (Figure 1C). Unexpectedly, the H3K27me3 histone mark, which is deposited by the polycomb repressive complex^38^, was neither correlated with positive nor with negative eigenvector coefficients **(Extended Data Figure 1B).** We then speculated that, as H3K27me3 was not related with standard A or B compartments, this histone mark may be linked to a different type of chromatin compartmentalization. In this context, a visual inspection of the first eigenvector distribution revealed a positive extreme, a negative extreme and a long intermediate valley (Figure 1F). Indeed, applying the Bayesian Information Criterion, we observed that a classification into three compartments was the best compromise between distribution fitting accuracy and minimum number of compartments **(Extended Data Figure 1D).** Subsequently, we modelled the eigenvector distribution to establish the thresholds segmenting the data into an A-type, B-type and intermediate (i)-type compartments **(Extended data Figure 1E** and Figure 1F). Analyzing these three compartments together with other omics layers revealed the expected association of A-type compartment with active chromatin, B-type compartment with H3K9me3, and a remarkably association between the I-type compartment and the presence of H3K27me3 (Figure 1G). Indeed, a chromHMM-based chromatin state model specific for B cells^28, 39^ revealed that the regions associated with the I-type compartment were enriched for poised-promoter and polycomb-repressed chromatin states (Figure 2A and **Extended Data Figure 2A).**

**Figure 2.**
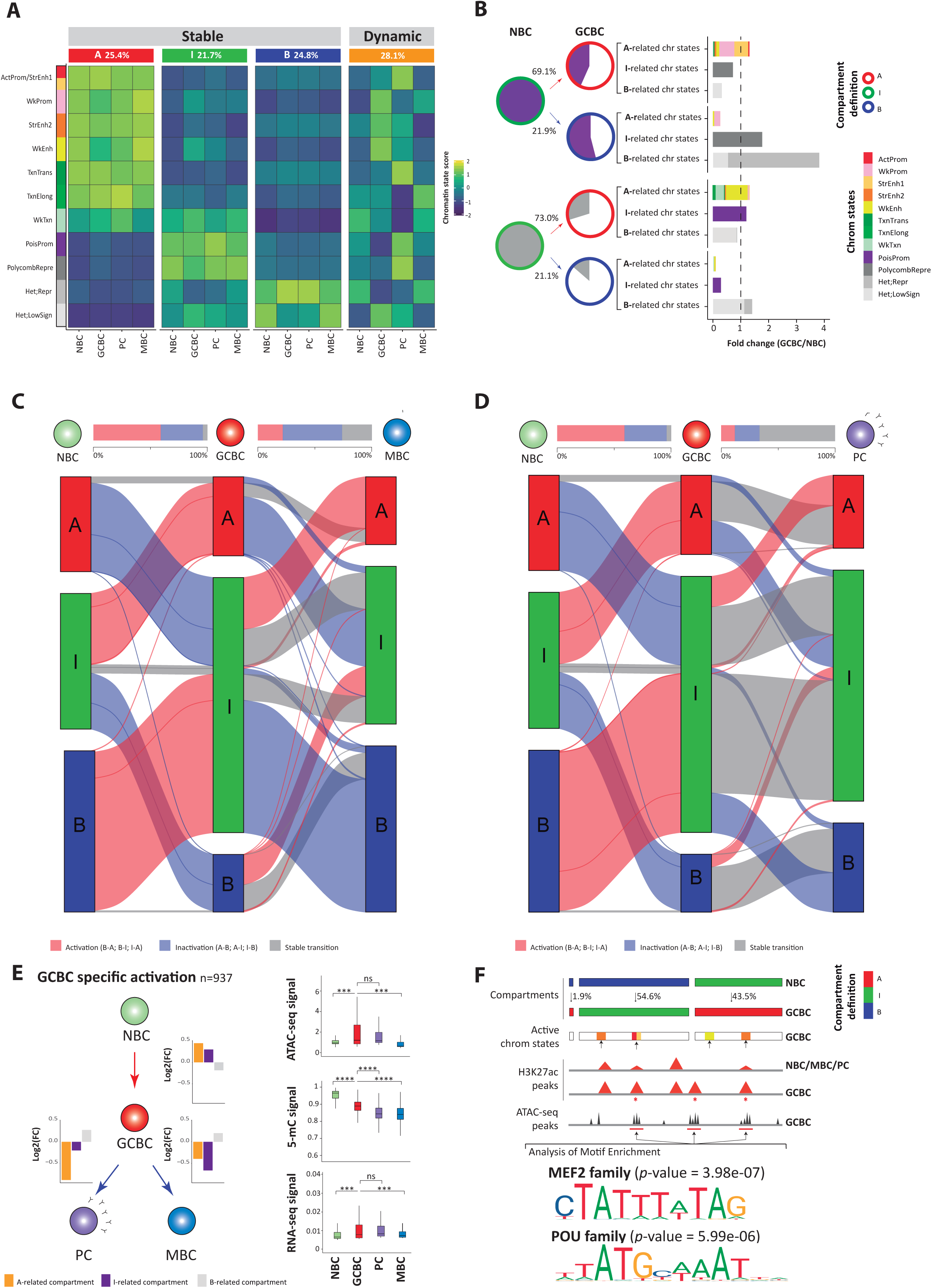
Chromatin dynamics across B cell differentiation. **A** - Functional association of the conserved and dynamic compartments during B cell maturation using eleven chromatin states (normalized by sample and chromatin state). Conserved compartments were segmented into A-type, I-type and B-type compartments. The percentage of each conserved or dynamic compartment is indicated for all B cell subpopulations. ActProm-StrEnhl, Active Promoter-Strong Enhancer 1; WkProm, Weak Promoter; StrEnh2, Strong Enhancer 2; WkEnh, Weak Enhancer; TxnTrans, Transcription Transition; TxnElong, Transcription Elongation; WkTxn, Weak Transcription; PoisProm, Poised promoter; Polycom bRepr, Polycomb repressed; Het;Repr, Heterochromatin;Repressed; Het;LowSign, Heterochromatin;Low Signal. **B -** Intermediate compartment dynamics. Pie charts represent poised promoters (top, violet color) or polycomb-repressed (bottom, light gray color) within the I-type compartment in NBC which shifts to A-type and B-type compartments in GCBC. The pie charts under GCBC represent the fraction that maintains the previous chromatin state (colored as previously defined) or changed chromatin states (not colored). Bar graphs represent the fold change between GCBC and NBC of each three groups of chromatin states (arranged by their relationship to the A-type, I-type and B-type compartments). Active Promoter, Weak Promoter, Strong Enhancer 1, Strong Enhancer 2, Weak Enhancer, Transcription Transition, Transcription Elongation, Weak Transcription were A-type compartment-related states. Heterochromatin/Repressed and Heterochromatin/Low signal were B-type compartment-related states. Poised Promoter or Polycomb repressed chromatin states were I-type compartment-related states. **C/D-** Alluvial diagrams showing the compartment dynamics in the two branches of mature B cell differentiation: NBC-GCBC-MBC **(C)** and NBC-GCBC-PC **(D).** Activation, in red, represents changes from compartment B-type to A-type, B-type to I-type and I-type to A-type. Inactivation, in blue, represents changes from A-type to B-type, A-type to I-type and I-type to B-type compartments. The non-changed compartments are represented in gray. On the top, the bar plots between B cell subpopulations represent the total percentage of regions changing to active or inactive, and regions that conserve its previous compartment definition. **E -** Multi-omics characterization of the 937 regions (of 100Kb resolution) gaining activity exclusively in GCBC. **Left:** Scheme of B cell differentiation and chromatin state dynamics, in which the barplots indicate the Iog2 fold change of active, intermediate or inactive -related chromatin state groups. **Right:** Boxplots of chromatin accessibility (ATAC-seq signal), DNA methylation (5-mC signal) and gene expression (RNA-seq signal) per B cell subpopulations compared using the Wilcoxon’s test. *p-value<0.05, **p-value<0.001, ***p-value<0.0001, ****p-value<0.00001. **F -** Enrichment analysis of transcription factor binding motifs. **Top:** Schematic representation of the analytic strategy. **Bottom:** Binding motifs of MEF2 and POU TF families are highly enriched in active and accessible loci in the GCBC specific regions gaining activity (n=171 independent genomic loci) versus the background (n=268 independent genomic loci), p-values were calculated using the AME-MEME suite. Out of the list of all enriched transcription factor binding motifs, we considered only those expressed in the three GCBC replicates.

We next quantified the compartment interactions by computing the compartment score (C-score) as the ratio of intra-compartment interactions over the total chromosomal interactions per compartment **(Extended Data Figure 2B).** Interestingly, the I-type compartment was associated with lower C-score than the A-type and B-type compartments **(Extended Data Figure 2C).** We further explored this phenomenon by dividing the I-type compartment into two blocks differentiating positive (IA) and negative (IB) eigenvector components **(Extended Data Figure 2D).** The analysis showed that the I-type compartment, regardless being IA or IB, was consistently having lower C-score than the A or B-type compartments. This finding further supports the existence of the I-type compartment as an independent chromatin structure different from A and B-type compartments. Additionally, it suggests that the I-type compartment tends to interact not only with itself but also with A and B-type compartments, and as such it may represent an interconnected space between the fully active and inactive compartments.

To study the potential role of the I-type compartment during B cell differentiation, we selected poised promoters or polycomb repressed regions within this compartment in NBC and studied how they change in both compartment and chromatin state upon differentiation into GCBC (Figure 2B). The majority of compartment transitions (69.1% of poised promoter and 73.0% of polycomb repressed) change into A-type compartment, a consistent fraction (21.9% and 21.1%) into B-type, and only a small fraction (9% and 5.9%) maintain their intermediate definition. This finding indicates that the regions with a most prominent I-type compartment character undergo a widespread structural modulation during NBC to GCBC differentiation step. Interestingly, transitions from I-type to A-type compartment (activation events) were paired with a reduction of poised promoters (56.7% loss) and polycomb repressed states (70.2% loss). These reductions were associated with an increase of A-related chromatin states (1.31- or 1.33-fold change coming from poised promoter or polycomb-repressed, respectively) such as promoter, enhancer and transcription (Figure 2B). Conversely, poised promoters and polycomb-repressed regions associated with I-type compartments in NBC that changed into B compartments in GCBC (inactivation events) were related to an increase of B-related chromatin states (3.81 or 1.4-fold change coming from poised promoter or polycomb-repressed, respectively) such as heterochromatin characterized by H3K9me3 (Figure 2B).

Altogether, these results point to the existence of an intermediate transitional compartment with biological significance, enriched in poised and polycomb-repressed chromatin states, interconnected with A and B -type compartments, and amenable to rewire the pattern of interactions leading to active or inactive chromatin state transitions upon cell differentiation.

### Changes in genome compartmentalization are reversible during B cell differentiation

Mapping A, I and B-type compartments in NBC, GCBC, MBC and PC Hi-C maps revealed that 28.1% of the genome dynamically changes compartment during B cell differentiation (Figure 2A and **Extended Data Figure 2A).** B cell differentiation is not a linear process, NBC differentiate into GCBC, which then branch into long-lived MBC or antibody-producing PC. Thus, we studied the 3D genome compartment dynamics along these two main differentiation paths (NBC-GCBC-PC and NBC-GCBC-MBC). At each differentiation step, we classified the genome into three different dynamics: (i) compartments undergoing activation events (B-type to A-type, B-type to I-type, or I-type to A-type), (ii) compartments undergoing inactivation events (A-type to B-type, A-type to I-type, or I-type to B-type), and (iii) stable compartments (Figure 2C-D). The NBC-GCBC-MBC differentiation path suggests that the extensive remodeling taking place from NBC to GCBC is followed by an overall reversion of the compartmentalization in MBC, achieving a profile similar to NBC (Figure 2C). To assess the capacity of the genome to revert to a past 3D configuration, we analyzed the compartments in NBC as compared to those in PC and MBC. Indeed, we globally observed that 72.7% of the regions in MBC re-acquire the same compartment type as in NBC. This phenomenon was mostly related to compartments undergoing activation in GCBC, as 82.9% of them reverted to inactivation upon differentiation into MBC. This finding is in line with solid evidence showing that NBC and MBC, in spite of representing markedly different maturation B cell stages, are phenotypically similar^40, 41^ (Figure 1D). In the case of PC, the compartment reversibility accounted only for 30.8% of the genome (Figure 2D). To determine whether this compartment reversibility was also accompanied by a functional change, we analyzed the chromatin state dynamics within the compartments becoming uniquely active in GCBC as compared to NBC, MBC and PC (n=937) (**Supplementary Table 2**). We observed that the transient compartment activation from NBC to GCBC is related to an increase of A-related chromatin states (1.36-fold change). Conversely, the subsequent 3D genome inactivation upon differentiation into MBC and PC was related to an increase in B-related chromatin states (1.21- and 1.15-fold change, respectively) (Figure 2E left). Furthermore, those regions had a significant increase in chromatin accessibility and gene expression in GCBC as compared to NBC and MBC, but not in PC (Figure 2E right). These findings suggest that structural 3D reversibility in MBC is accompanied by a functional reversibility whereas PC partially maintains gene expression levels and chromatin accessibility similar to GCBC in spite of the compartment changes. Interestingly, in contrast to chromatin-based marks, DNA methylation was overall unrelated to compartment or chromatin state dynamics of the B cell differentiation (Figure 2E right).

### The 3D genome of GCBC undergoes extensive compartment activation

Our analyses revealed that the NBC and GCBC transition was associated to a large structural reconfiguration of compartments involving 96.0% of all dynamic compartments (Figure 2A). Interestingly, 61.5% of the changes between NBC and GCBC involved compartment activation (Figure 2C-D). As the germinal center reaction is known to be mediated by specific transcription factors (TFs)^42,43^ and those may be involved in shaping the spatial organization of the genome^8,14,16^, we further explored the presence of TF binding motifs in the newly activated compartments. We identified significantly enriched motifs for MEF2 and POU families (Figure 2F and **Supplementary Table 3)**, which are essential TFs involved in germinal center formation^44–47^. Furthermore, the newly activated compartments hosted about 100 genes significantly upregulated in GCBC as compared to NBC (FDR<0.05) (**Supplementary Table 4**). Remarkably, among them was the Activation Induced Cytidine Deaminase *(AICDA)* gene, which is essential for class-switch recombination and somatic hypermutation in GCBC and is specifically expressed in GCBC^48^. Indeed, the *AICDA* locus was globally remodeled from an inactive state in NBC to a global chromatin activation in GCBC, which included an increase in the ratio of GCBC/NBC 3D interactions as well as increased levels of active chromatin states (that is, active promoter and enhancers as well as transcriptional elongation), open chromatin, and gene expression (Figure 3A-B). This analysis also revealed the presence of possible upstream and downstream AICDA-specific enhances that gain interactions with the gene promoter in GCBC (Figure 3B). Interestingly, this multilayer chromatin activation at the *AICDA* locus was reverted to the inactive ground state once GCBC differentiate into MBC or PC.

**Figure 3.**
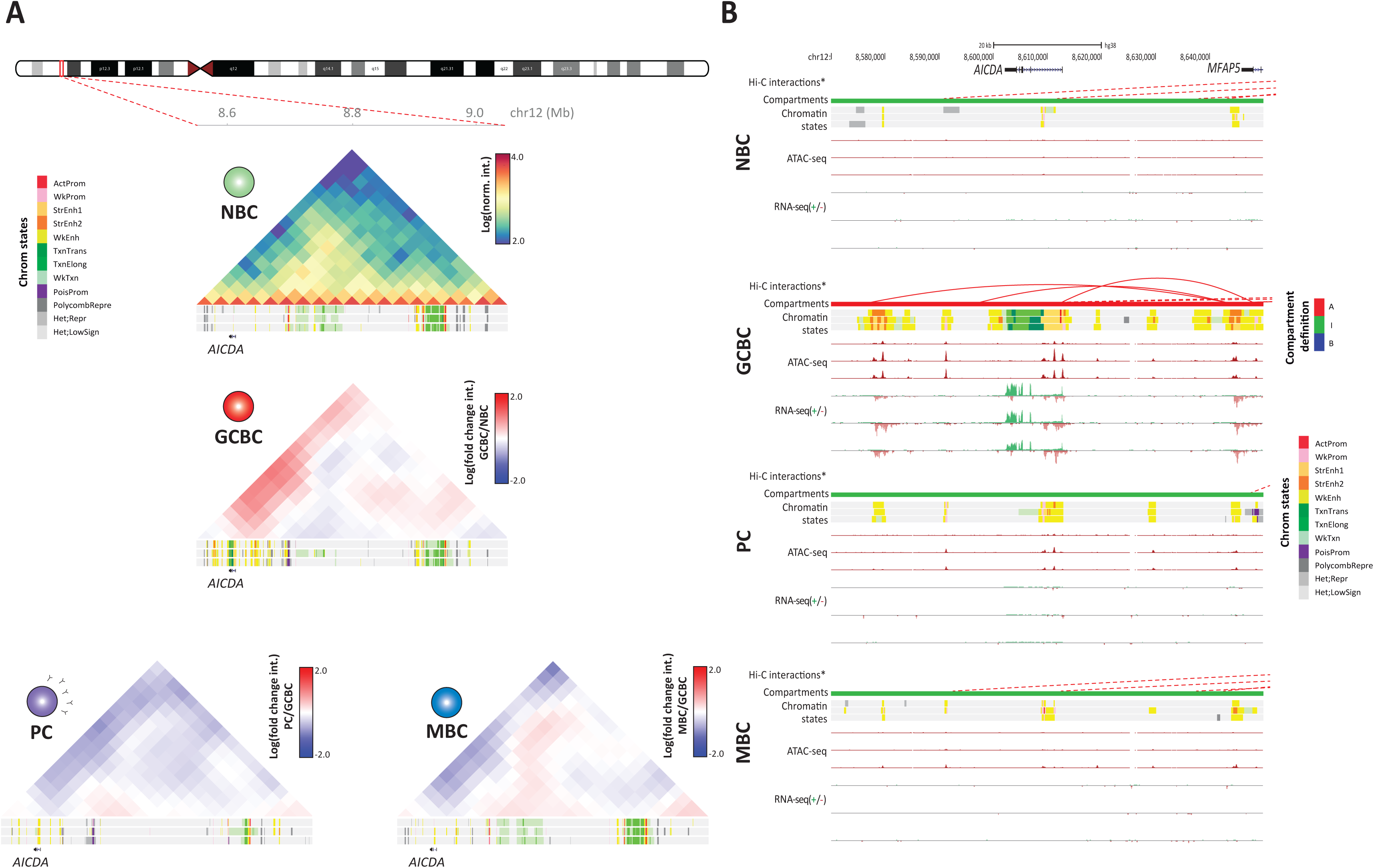
Chromatin organization at the *AICDA* locus. **A** - Normalized Hi-C contact map of the domain structure surrounding the *AICDA* gene in NBC. The log fold change interaction ratio between GCBC, MBC or PC as compared to NBC was computed. Below each interaction map, chromatin state tracks of three biological replicates per B cell subpopulation are shown. The coordinates of the represented region are chrl2:8,550,000-9,050,000, GRCh38. **B-** Multi-layer epigenomic characterization of *AICDA* gene region (chrl2:8,598,290-8,615,591, GRCh38) in four B cell subpopulations. Arc diagrams indicate the Hi-C significant interactions (continuous red lines involve the region of interest, while dashed red lines involve other regions of chromosome 12). Below them, we show compartment definition (red, compartment A-type: green, compartment I-type), chromatin states, chromatin accessibility (ATAC-seq, y-axis signal from 0 to 105) and gene expression (RNA-seq, y-axis signal from 0 to 4 for the positive strand and from 0 to −0.1 for the negative strand). Tracks of Hi-C interactions and compartment definition are based on merged replicates whereas chromatin states, chromatin accessibility and gene expression tracks of each replicate is shown separately. The coordinates of the represented region are chrl2:8,570,000-8,670,000, GRCh38.

### B cell neoplasms undergo disease-specific 3D genome reorganization

Next, we analyzed whether the observed 3D genome organization during normal B cell differentiation is further altered upon neoplastic transformation. To address this, we performed *in situ* Hi-C in fully characterized tumor cells from patients with chronic lymphocytic leukemia (CLL, n=7) or mantle cell lymphoma (MCL, n=5). Within each neoplasm, we included cases of two subtypes, IGVH mutated (m, n=5) and unmutated (u, n=2) CLL as well as conventional (c, n=2) and non-nodal leukemic (nn, n=3) MCL (Figure 4A and **Supplementary Table 5**). An initial unsupervised clustering of the RS from the entire Hi-C dataset indicated that CLL and MCL, similarly to the PCA from other omic layers generated from the same patient samples, clustered separately from each other and within a major cluster that included NBC and MBC (Figure 4B-C and **Extended Data Figure 4A).** Interestingly, NBC and MBC have been described as potential cells of origin of these neoplasms^22^. Furthermore, pairwise eigenvector correlation analysis of the cancer samples suggested that the 3D genome configuration of the two clinico-biological subtypes of CLL was rather homogeneous **(Extended Data Figure 4B-C).** This was not the case for the two MCL subtypes, which were more heterogeneous **(Extended Data Figure 4D-E).**

**Figure 4.**
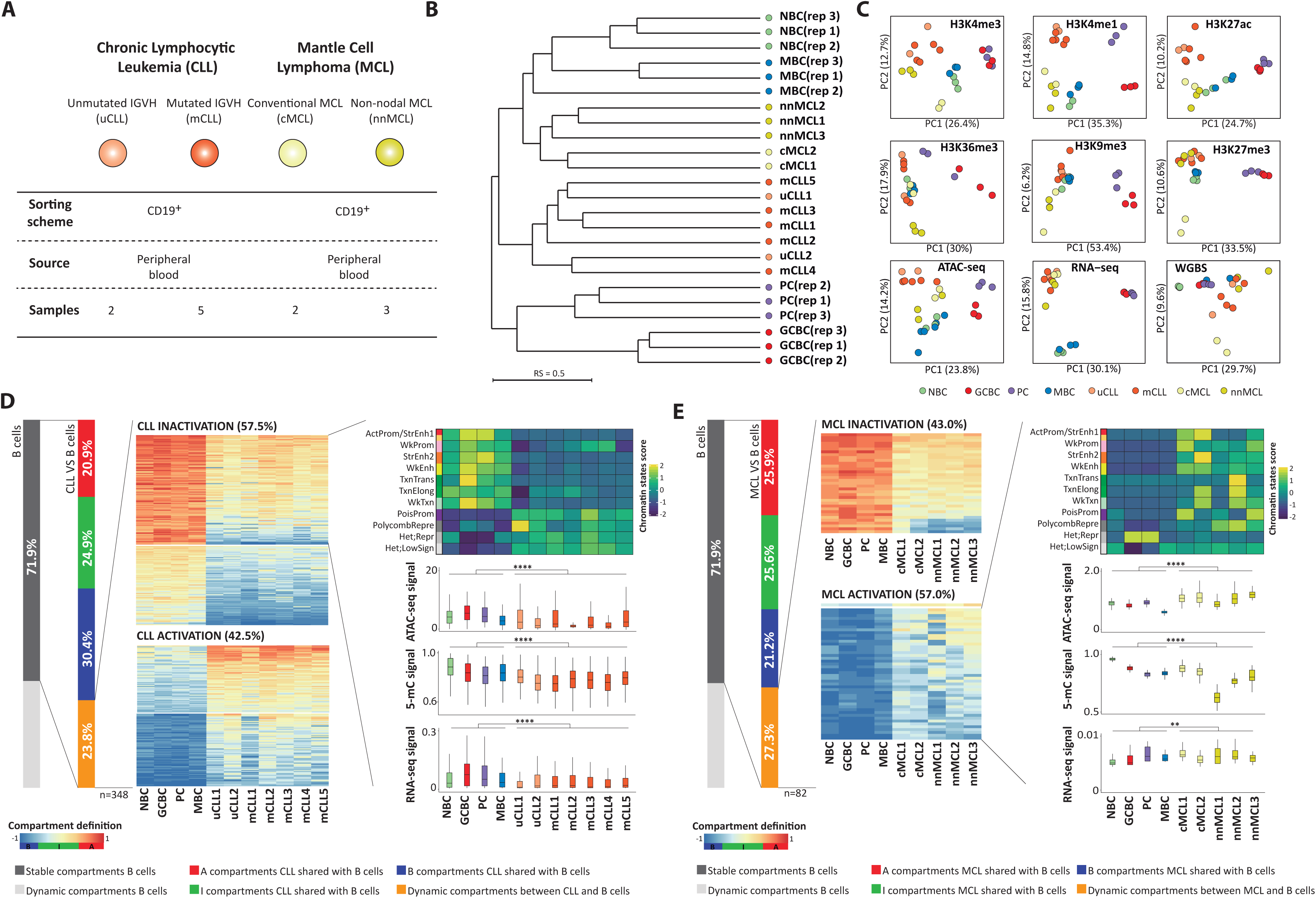
Characterization of the chromatin architecture of human B cell neoplasms. **A** - Sample description and *in situ* Hi-C experimental design in CLL and MCL cases. **B** - Dendrogram of the reproducibility score for normalized Hi-C contact maps at 100Kb for B cell subpopulations replicates and samples from B cell neoplasia patients. IGVH unmutated (u)CLL; IGVH mutated (m)CLL; conventional (c)MCLand non-nodal (nn)MCL. **C -** Unsupervised principal component analysis (PCA) for nine omic layers generated in the same patient samples as Hi-C: chromatin immunoprecipitation followed by sequencing (ChIP-seq) of six histone marks (H3K4me3 n=53,241 genomic regions, H3K4me1 n=54,653 genomic regions, H3K27Ac n=106,457 genomic regions, H3K36me3 n=50,530 genomic regions, H3K9me3 n=137,933 genomic regions, and H3K27me3 n=117,560 genomic regions), chromatin accessibility measured by ATAC-seq (n=140,187 genomic regions), DNA methylation measured by whole-genome bisulfite sequencing (WGBS, n=14,088,025 CpGs) and gene expression measured by RNA-seq (n=57,376 transcripts). In addition to the normal B cell subpopulations explained in Figure 1D, we studied 7 CLL patient samples (2 uCLL and 5 mCLL) and 5 MCL patient samples (2 cMCL and 3 nnMCL). **D -** Compartment changes upon CLL transformation. **Left:** First bar graph represents the percentage of conserved and dynamic compartments during normal B cell differentiation. Second bar graph shows the percentage of compartments stable and differential in CLL as compared to normal B cells. A total of 23.8% of the compartments change in at least one CLL sample. **Middle:** Heatmaps showing eigenvector coefficients of the 348 compartments significantly losing (n=200) or gaining activation (n=148) between all CLL samples and normal B cells. **Right:** Multi-omics characterization of the 200 regions losing activity in CLL. We show chromatin states, chromatin accessibility (ATAC-seq signal), DNA methylation (5-mC signal) and gene expression (RNA-seq signal) in CLL and normal B cells. Comparisons were performed using the Wilcoxon’s test. ****p-value<0.00001. **E -** Compartment changes upon MCL transformation. **Left:** First bar graph represents the percentage of conserved and dynamic compartments in B cells. Second bar graph shows the percentage of conserved compartments between B cells and MCL, being 27.29% non­conserved compartment in MCL. **Middle:** Heatmaps showing eigenvector coefficients of significant dynamic compartments (n=82) between MCL and B cells. Regions were split in two groups (MCL activation, n=35 or inactivation, n=47) according to the structural modulation of the MCL compared to B cells. **Right:** Example of the MCL activation subset (mostly those B-type compartments in B cells which significantly increase eigenvector coefficients in MCL) showing the chromatin states pattern, chromatin accessibility (ATAC-seq signal), DNA methylation (5-mC signal) and gene expression (RNA-seq signal). Comparisons were performed using the Wilcoxon’s test. *p-value<0.05, **p-value<0.001, ***p-value<0.0001, ****p-value<0.00001.

The differential clustering of CLL and MCL samples hint into disease-specific changes of their 3D genome organization (Figure 4B). To further detect those changes, we took the fraction of the genome with stable compartments during normal B cell differentiation and compared them to each lymphoid neoplasm. Qualitatively, we observed that roughly one quarter of the genome changes compartments in at least one CLL (23.8%) and at least one MCL sample (27.3%) as compared to normal B cells (Figure 4D-E left). Using a more stringent quantitative approach, we aimed at detecting changes associated with CLL or MCL as whole, which revealed a total of 348 and 82 significant compartment changes (absolute difference in the eigenvalue>0.4 and FDR<0.05) in CLL and MCL, respectively. The larger number of regions changing compartments in CLL correlates with the results of the Hi-C based clustering (Figure 4B), which indicates that MCL is more similar to NBC/MBC than CLL. Moreover, the observed compartment changes tended towards inactivation in CLL (57.5%) (Figure 4D middle) and towards activation in MCL (57.0%) (Figure 4E middle) compared to the normal B cells. These 3D genome organization changes were associated with the expected changes in chromatin function. Inactivation at the 3D genome level in CLL was linked to a shift to poised promoter and polycomb-repressed chromatin states, and a significant loss of chromatin accessibility and gene expression (Figure 4D right). Activation at the 3D genome level in MCL was accompanied with an enrichment of active chromatin states and a significantly increase in chromatin accessibility and gene expression (Figure 4E right). Overall, these results point to the presence of recurrent and specific changes in the 3D genome organization in CLL and MCL, being the former more extensively altered than the latter.

### *EBF1* downregulation in CLL is linked to extensive 3D genome reorganization

To further characterize the compartmentalization of neoplastic B cells, we classified the changing compartments as common (between CLL and MCL) or entity-specific (either in CLL or MCL). We detected 31 compartments commonly altered in both malignancies, revealing the existence of a core of regions that distinguish normal and neoplastic B cells (Figure 5A-B). A targeted analysis of CLL and MCL revealed 89 CLL-specific (41 and 48 inactivated and activated, respectively) and only 3 MCL-specific compartment changes (Figure 5C, Figure 6A and **Extended Data Figure 5A).** Interestingly, the set of 41 compartments inactivated in CLL were significantly enriched (p-value=0.0060) in downregulated genes (n=ll) as compared to normal B cells and MCL samples, being the Early B cell Factor 1 *(EBF1)* a remarkable example (Figure 5C-D and **Supplementary Table 6**). *EBF1* downregulation has been described to be a diagnostic marker in CLL^49^, and its low expression may lead to reduced levels of numerous B cell signaling factors contributing to the anergic signature of CLL cells^50, 51^ and low susceptibility to host immunorecognition^52, 53^. To obtain insights into the mechanisms underlying *EBF1* silencing in CLL, we analyzed in detail a 2Mb region hosting the gene, which also contains two nearby protein coding genes, *RNF145* and *UBLCP1,* and a IncRNA, *LINC02202. We* observed that a large fraction of 3D interactions involving the *EBF1* region in normal B cells were lost in CLL resulting in a change from A-type to I-type compartment and a sharp inactivation of the gene, as shown by the analysis of chromatin states (Figure 5E). Remarkably, in spite of the global reduction of 3D interactions, the two adjacent genes *(RNF145* and *UBLCP1)* were located in the only region (spanning 200Kb) that remained as A-type compartment in the entire 2Mb region, maintaining thus an active state. To obtain further insights into the *EBF1* genome structure, we modeled its spatial organization in NBC and CLL by using the restraint-based modeling approach implemented in TADbit^54, 55^ (Figure 5F and **Extended Data Figure 5B-C).** The *EBF1* domain in CLL resulted in larger structural variability as compared with the models in NBC due to the depletion of interactions in neoplastic cells **(Extended Data Figure 5B).** The 3D models revealed that the *EBF1* gene is located in a topological domain, isolated from the rest of the region in NBC, hosting active enhancer elements (Figure 5F). Remarkably, the active enhancer elements together with the interactions are lost in CLL (Figure 5F), resulting in more collapsed conformations (Figure 5G). Overall, these analyses suggest that *EBF1* silencing in CLL is linked to a compartment shift of a large genomic region leading to the abrogation of interactions and regulatory elements.

**Figure 5.**
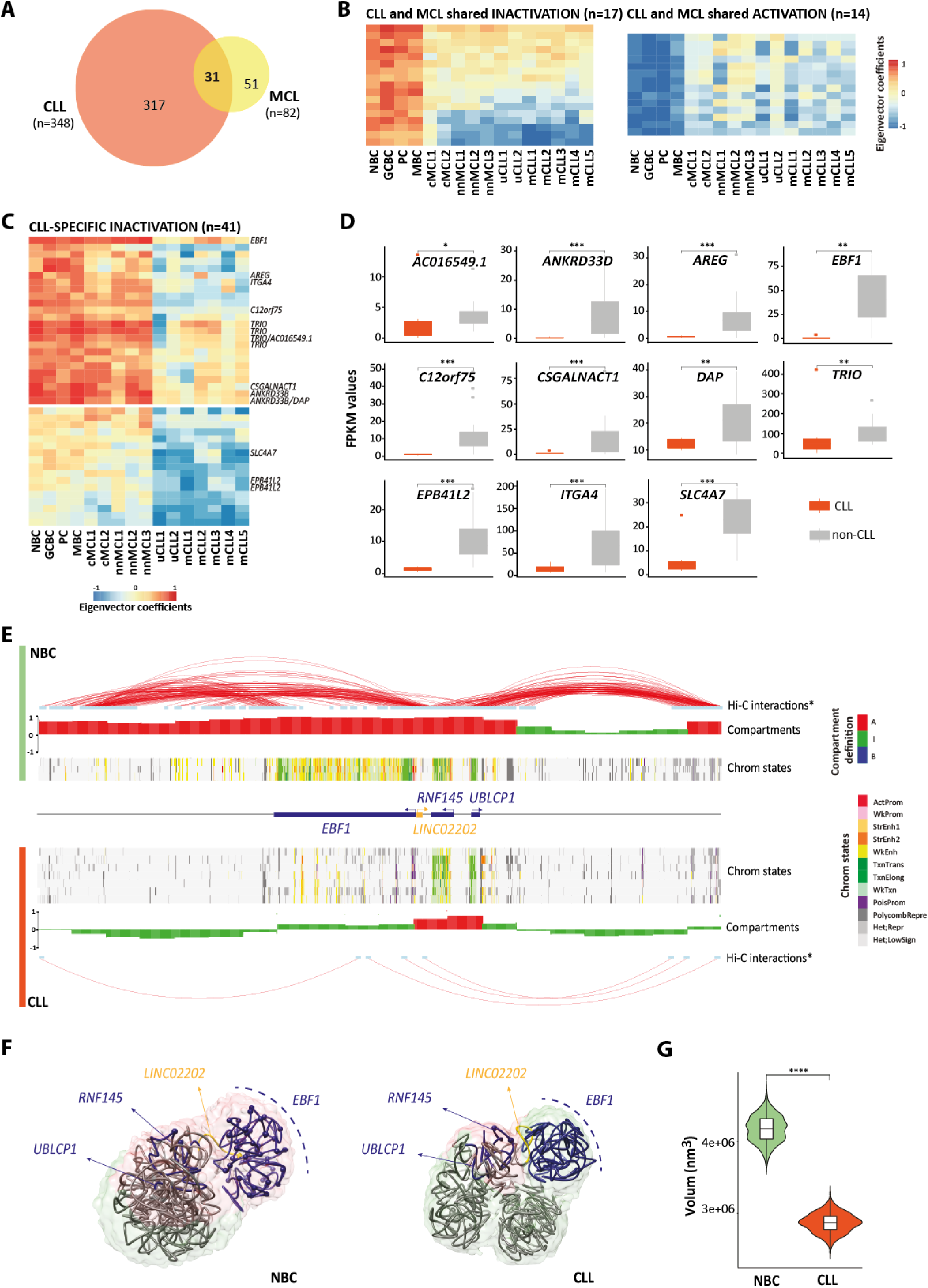
*EBF1* silencing in CLL is accompanied by structural changes affecting a 2Mb region. **A** - Venn diagram showing the significant number of dynamic compartments in CLL and MCL as compared to normal B cell differentiation and the regions shared between both B cell neoplasms (n=31). **B -** Heatmaps showing eigenvector coefficients of compartments significantly losing or gaining activation between B cell neoplasms (MCL and CLL together) and B cells. **C -** Heatmap showing the eigenvector coefficients of the compartments losing activation specifically in CLL (n=41). Significantly downregulated genes (FDR<0.05) associated to each compartment are shown on the right of the heatmap (p-value=0.0038, calculated from the total number of genes picked on 48 random compartments per 10,000 times). **D -** FPKM values of all the CLL-specific significantly downregulated genes within compartments losing activation. *adjusted p-value<0.05, **adjusted p-value<0.005, ***adjusted *p-* value<0.0005. **E -** Map of the *EBF1* regulatory landscape. Significant Hi-C interactions (p-value=0.001) and compartment type from merged NBC and a representative CLL sample, followed by chromatin state tracks from each NBC (n=3) and CLL (n=7). The coordinates of the represented region are chr5:158,000,000-160,000,000, GRCh38. **F -** Restraint-based model at 5Kb resolution of the 2Mb region containing *EBF1* (total 400 particles, *EBF1* locus localized from 139 to 220 particle). Data from merged NBC **(top)** and CLL **(bottom)** was used. Surface represents the ensemble of 1,000 models and is color-coded based on the compartment definition (A-type, B-type and I-type in red, blue and green, respectively). The top-scoring model is shown as trace, where protein-coding genes are colored in blue and long non-coding RNAs in yellow. Spheres represents enhancer regions. **G -** Violinplot of the convex hull volume involving the 81 particles from the *EBF1* region. Comparison was performed using Wilcoxon’s test. p-va I ue=0.00001.

**Figure 6.**
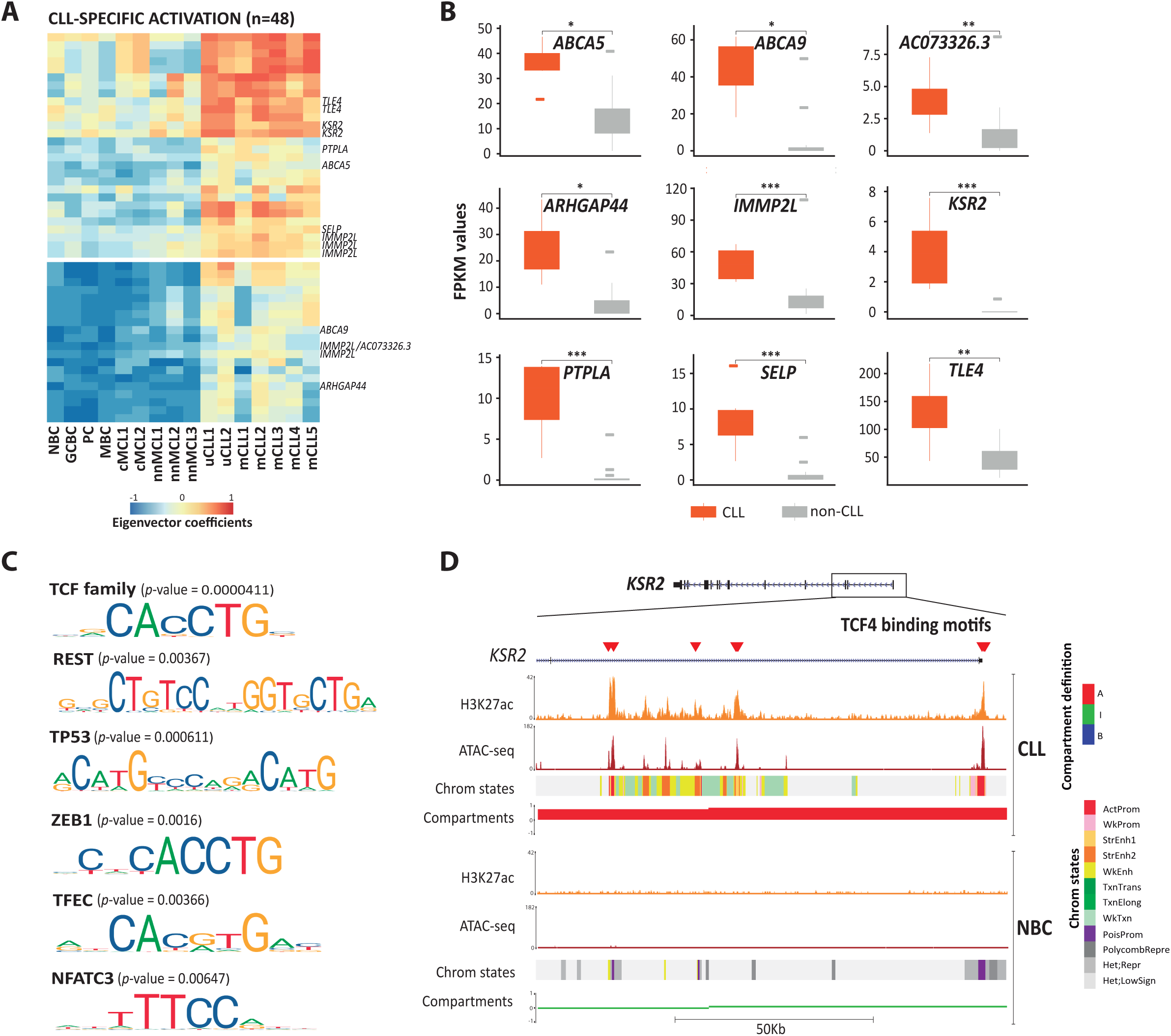
Transcription factors associated to CLL-specific activated compartments. **A** - Heatmap showing the first eigenvector coefficients of the compartments gaining activation specifically in CLL (n=48). Significantly upregulated genes (FDR=0.05) associated to each compartment are shown on the right of the heatmap (p-value=0.006). **B -** FPKM values of all the CLL-specific significantly upregulated genes within compartments gaining activation. *adjusted p-value<0.05, **adjusted p-value<0.005, ***adjusted *p-* value<0.0005. **C -** Enrichment analysis of transcription factor binding motifs. We show the most significant TF binding motifs enriched in active and accessible loci within the CLL-specific regions gaining activity (n=25 independent genomic loci) versus the background (n=28 independent genomic loci), p-values were calculated using the AME-MEME suite. Out of the list of all enriched transcription factor binding motifs, we considered only those expressed in all CLL samples (n=7). **D -** Example of TCF4 binding motifs at the *KSR2* promoter region in CLL and NBC. We show the following tracks: H3K27ac, chromatin accessibility (ATAC-seq) and chromatin states of a representative NBC replicate and CLL sample. The coordinates of the represented region are chrl2:117,856,977-117,975,164, GRCh38.

Our analysis also detected 48 regions that changed towards more active compartment exclusively in CLL (Figure 6A). As expected, these regions were significantly enriched in upregulated genes (p-value=0.0038) and harbored 9 genes with increased expression (Figure 6B and **Supplementary Table 7**). As previously shown for regions gaining activity in GCBC (Figure 2E), we evaluated whether particular TFs were related to the CLL-specific increase in 3D interactions. Indeed, we found an enrichment in TF binding motifs of the TCF (p-value=0.00004) and NFAT (p-value=0.00647) families, which have been described to be relevant for CLL pathogenesis^28, 56, 57^ (Figure 6C and **Supplementary Table 8**). One of the nine upregulated genes in CLL-specific active compartments was *KSR2,* a gene whose upregulation has a strong diagnostic value in CLL^49^. Importantly, this gene contained several motifs for the TCF4 transcription factor (Figure 6D), which itself is overexpressed in CLL as compared to normal B cells^28^, suggesting in this particular example that TCF4 overexpression may lead to aberrant binding to *KSR2* regulatory elements and a global remodeling of its 3D interactions.

### Increased 3D interactions across a 6.1Mb region including the 50X11 oncogene in aggressive MCL

In addition to entity-specific 3D genome changes, our initial analyses also suggested that different clinico-biological subtypes may have a different 3D genome organization, especially in MCL (Figure 4B). To identify subtype differences within each B cell neoplasia, we selected regions with homogeneous compartments within each disease subtype and classified them as distinct if the difference between the Hi-C matrices cross-correlation eigenvalues was greater than 0.4. Applying this criterion, we defined 47 compartment changes between uCLL and mCLL, and 673 compartment changes between nnMCL and cMCL (Figure 7A). This finding confirmed the previous analyses **(Extended Data Figure 4B-E),** and indicated that the two MCL subtypes have a markedly different 3D genome organization. Two thirds of the compartments changing in the MCL subtypes (n=435, 64.6%) gained activity in the clinically aggressive cMCL, and one third gained activity in nnMCL. We then characterized the chromosomal distribution of these compartment shifts, which, surprisingly, was significantly biased towards specific chromosomes (Figure 7B). In particular, those regions gaining 3D interactions in aggressive cMCL were highly enriched in chromosome 2, being 22.3% (n=97) of all 100Kb compartments located in that chromosome (Figure 7B). We next analyzed chromosome 2 of cMCL in detail and we observed a *de novo* gain of A-type and I-type compartments accumulated at band 2p25 as compared to both normal B cells and nnMCL (Figure 7C). The entire region of about 6.1Mb had a dramatic increase of interactions and active chromatin states in cMCL as compared to nnMCL (Figure 7D and **Extended Data Figure 7A).** Most interestingly, this region contains *SOX11,* whose overexpression in cMCL represents the main molecular marker to differentiate these two MCL subtypes^58^, and has been shown to play multiple oncogenic functions in cMCL pathogenesis^59–61^. However, as *SOX11* is embedded into a large block of 6.1Mb gaining activation in cMCL, we wondered whether additional genes could also become upregulated as a consequence of the large-scale spatial organization of chromosomal band 2p25. Indeed, mining the expression data from the 5 MCL cases studied herein as well as two additional published cohorts^49, 62^, we observed that 13 (43%) of the 30 expressed genes within the 6.1Mb region were over-expressed in cMCL as compared to nnMCL in at least one cohort (Figure 7D and **Extended Data Figure 7B-C),** which may also contribute to cMCL pathogenesis and clinical aggressiveness.

**Figure 7.**
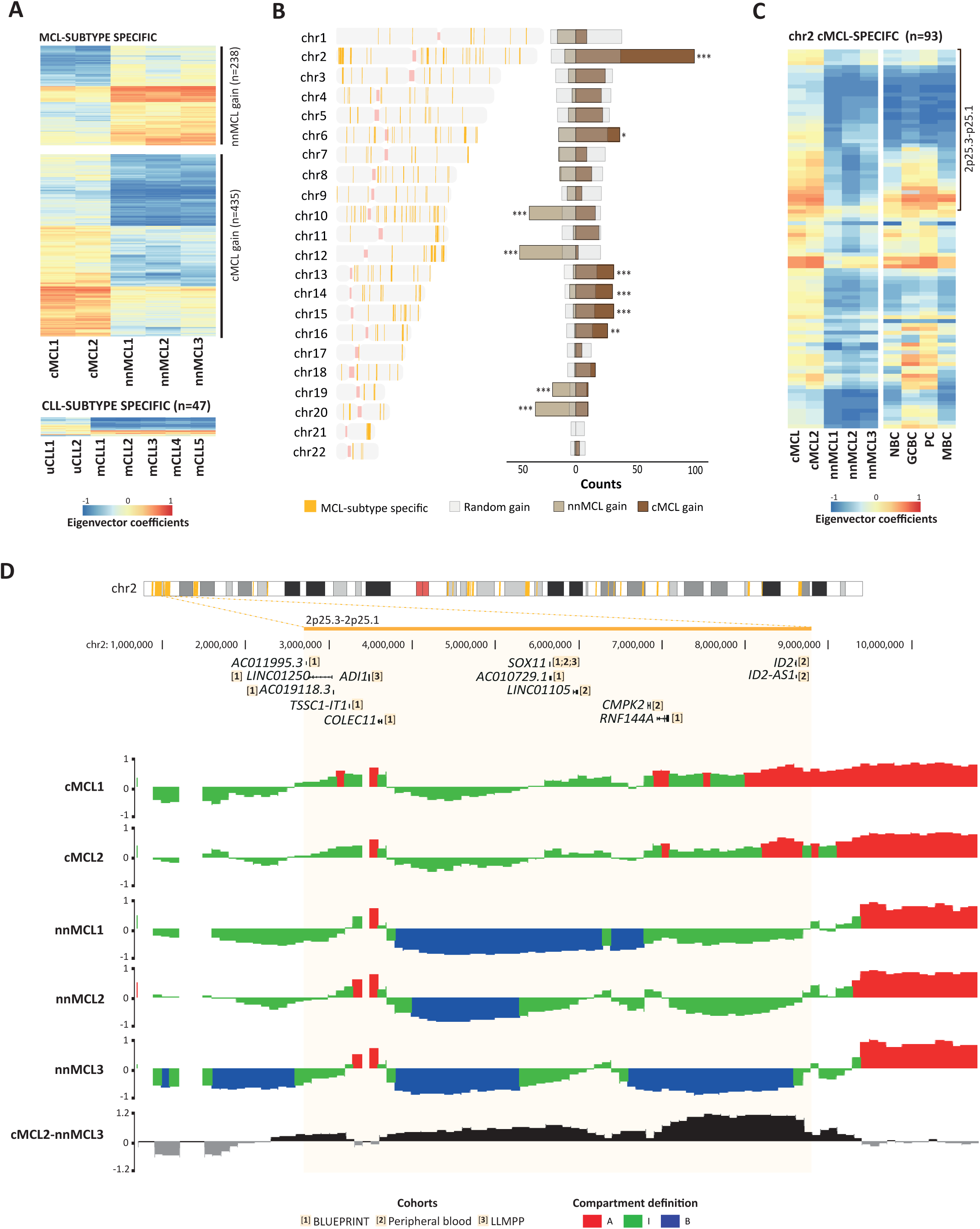
Long-range chromatin remodeling of a 6.1Mb involving 50X11 in cMCL. **A** - Heatmaps showing eigenvector coefficients of compartments significantly changing in cMCL versus nnMCL (n=673) and in uCLL versus mCLL (n=47). **B - Left:** Genome-wide distribution of compartments changing in MCL subtypes. The vertical orange lines point to the chromosome location of the regions. **Right:** Relative abundance of the compartments significantly gaining activity in cMCL or nnMCL as compared with a random probability. A gain in compartment activation was defined as an increase of eigenvector coefficient of at least 0.4. *p-value<0.05, **p-value<0.005, ***p-value<0.0005. **C -** Heatmap showing eigenvector coefficients of the chromosome 2 compartments specifically gaining activation in cMCL (n=93). On the top of the heatmap, we show the 6.1Mb genomic block gaining activation in 2p25. **D - Top:** Differentially expressed genes between cMCL and nnMCL in each of the three cohorts of transcriptional data of MCL patients. **Bottom:** Compartment type tracks on all the MCL samples under study. Eigenvalue subtraction between representative cMCL and nnMCL samples highlighting the 6.1Mb region gaining activity in the former.

## Discussion

We present a comprehensive analysis of the dynamic genome architecture reorganization during normal human B cell differentiation and upon neoplastic transformation into CLL and MCL. The integration of 3D genome data with nine additional omics layers including DNA methylation, chromatin accessibility, six histone modifications and gene expression, has allowed us to obtain new insights into 3D genome functional compartmentalization, cellular transitions across B cell differentiation and 3D genome aberrations in neoplastic B cells. We initially explored the distribution of Hi-C eigenvector coefficients and identified that a categorization into three components seemed to be more appropriate than the well-established dichotomous separation of the genome into A and B compartments^5^. Between the active (A) and repressed (B) compartments, we revealed the presence of an intermediate (I) component which contained more inter-compartment interactions than fully active or inactive chromatin, and is enriched in H3K27me3 located within poised promoters and polycomb-repressive chromatin states. Thus, this I-type compartment may represent a labile state of the high-order chromatin organization that may evolve either into active or inactive chromatin compartments. The existence of an intermediate compartment may be supported by several lines of published evidence. For example, during T cell commitment, a correlation between intermediate compartment scores with intermediate levels of gene expression was observed^15^. Recently, using super-resolution imaging, it was found that some compartments could belong to active or inactive states depending on the observed cell^63^, which could resemble an intermediate compartment in a population-based analysis such as Hi-C. Finally, these evidences are also in line with the observation that the polycomb repressive complex forms discrete subnuclear chromatin domains^64–66^ that can be dynamically modulated during cell differentiation^67, 68^.

The three compartments had extensive modulation during human B cell differentiation, a process whose 3D genome architecture has been previously studied in cell lines and primary mouse cells^8,14,69–72^ or during the human germinal center reaction^16^. We observed that 28.1% of the genome is dynamically altered in particular B cell maturation transitions, a magnitude that is in line with compartment transitions observed during the differentiation of human embryonic stem cells into four cell lineages^7^ or the reprogramming of mouse somatic cells into induced pluripotent stem cells^8, 73^, but lower than an analysis of compartment transitions across 21 human cells and tissues, which reached 60% of the genome^13^. The compartment modulation linked to B cell maturation was mainly related to two phenomena, a large-scale activation from NCB to GCBC and a reversion of the 3D genome organization of MBC back to the one observed in less mature NBC. As the number of mid-range 3D interactions upon activation has been suggested to decrease^74^, our result on the GCBC structural activation supports a previous study in which the chromatin structure of GCBC undergoes global de-compaction^16^. In this context, TFs have been described to act as the architects instructing structural changes in the genome^75^ and a recent report has described that TFs are able to drive topological genome reorganizations even before detectable changes in gene expression^8^. A detailed analysis of regions that become exclusively active in GCBC as compared to any other B cell subpopulation under study revealed an enrichment in TF binding motifs of MEF2 and POU families, which have been described to play a key role in the germinal center formation^43^. In line with this important role of TFs in activating chromatin in GCBC, we also identified that

NFAT and TCF binding motifs are enriched in those compartments specifically activated in CLL, and these TFs have also been previously linked to *de novo* active regulatory elements in CLL and its pathobiology^28^. All these results are concordant with studies in which lineage-restricted transcription factors have been proposed to establish and maintain genome architecture of specific lineages^14, 75–77^. The outcome of the germinal center reaction are PC and MBC, which are phenotypically and functionally distinct subpopulations. GCBC and PC show an overall high level of conservation of their 3D genome organization, but the differentiation into MBC is related to extensive changes. Remarkably, we observed roughly three quarters of the changes in MBC compartments reverted back to the compartment profile observed in NBC. This reversibility of the higher-order chromatin structure is very much in line with the previously observed similarity of histone modifications, chromatin accessibility and gene expression profiles in NBC and MBC. In sharp contrast to this congruent behavior of chromatin-based traits, DNA methylation is rather different between NBC and MBC, as this mark follows an accumulative pattern during cell differentiation^30, 78^ and can be used to faithfully track the lineage trajectory of the cells^79^.

We describe that B cell neoplasms show tumor-specific changes in the 3D genome organization that can span over large DNA stretches and contain genes linked to their pathogenesis. Of particular interest was the observation of the structural activation of 6.1Mb affecting the entire chromosome band 2p25.2 in aggressive cMCL, which contains the *SOX11* oncogene, a biomarker whose expression defines this MCL subtype^58^ and plays key functional roles in its pathogenesis^80^. Although the *SOX11* oncogene expression is related to the presence of active histone modifications in the promoter region^81^ and the establishment of novel 3D loops with a distant enhancer element^32^, our finding indicates that such looping is embedded into long-range alterations in the 3D genome structure. This change is not only linked to *SOX11* overexpression, but seems to be related to the simultaneous overexpression of multiple genes within the target region. This phenomenon of long-range epigenetic changes has been observed at the DNA methylation level, as the hypermethylation over one chromosomal band of 4Mb that has been linked to silencing of several genes in colorectal cancer^82^. Additionally, in prostate cancer, long-range chromatin activation or inactivation analyzed by histone modifications has been shown to target oncogenes, microRNAs and cancer biomarker genes^83^. The presence of large-range epigenetic remodeling in cancer^82–91^ shall support a more generalized use of genome-wide chromosome conformation capture techniques as part of the global characterization of primary human tumors. Beyond the identification of a concerted deregulation of multiple contiguous genes with a potential role in cancer biology, targeting long-range aberrations in the 3D genome structure may itself represent a therapeutic target.

In conclusion, we provide an integrative and functional view of the 3D genome topology during human B cell differentiation and neoplastic transformation. Beyond revealing the presence of a novel compartment related to the polycomb-repressive complex, our analysis points to a highly dynamic 3D genome organization in normal B cells, including extensive activation from NBC to GCBC and a reversibility in MBC. In neoplastic cells from CLL and MCL, we identify disease and subtype-specific change in the 3D genome organization, which include large chromatin blocks containing genes playing key roles in their pathogenesis and clinical behavior.

## Acknowledgments

This research was funded by the European Union’s Seventh Framework Programme through the Blueprint Consortium (grant agreement 282510), the World Wide Cancer Research Foundation Grant No. 16-1285 (to J.I.M-S.), the ERC (grant agreement 609989 to M.A.M-R.), European Union’s Horizon 2020 research and innovation programme (grant agreement 676556 to M.A.M-R.). We also knowledge the support of Spanish Ministerio de Ciencia, Innovación y Universidades through SAF2012-31138 and SAF2017-86126-R to J.I.M-S., SAF2015-64885-R to E.C., BFU2017-85926-P to M.A.M-R. and PMP15/00007 to E.C. which is part of Plan Nacional de l+D+l and co-financed by the ISCIII-Sub-Directorate General for Evaluation and the European Regional Development Fund (FEDER-“Una manera de Hacer Europa”) (to E.C.), the International Cancer Genome Consortium (Chronic Lymphocytic Leukemia Genome consortium to E.C.), La Caixa Foundation (CLLEvolution-HE17-00221, to E.C.). Furthermore, the authors would like to thank the support of the Generalitat de Catalunya Suport Grups de Recerca AGAUR 2017-SGR-736 (to J.I.M.-S.), 2017-SGR-1142 (to E.C.) and 2017-SGR-468 (to E.C.), the Accelerator award CRUK/AIRC/AECC joint funder-partnership, the CERCA Programme/Generalitat de Catalunya and CIBERONC (CB16/12/00225, CB16/12/00334 and CB16/12/00489). R.V-B. (BES-2013-064328) and P.S-V. (BES-2014-070327) were supported by a predoctoral FPI Fellowship from the Spanish Government and N.R. by the Acció instrumental d’incorporacio de científics i tecnòlegs PERIS 2016 from the Generalitat de Catalunya. The authors thank the Barcelona Supercomputing Center for access to computational resources. This work was partially developed at the Centro Esther Koplowitz (CEK, Barcelona, Spain). CRG acknowledges support from ‘Centro de Excelencia Severo Ochoa 2013-2017’, SEV-2012-0208 and the CERCA Programme/Generalitat de Catalunya. We thank Marta Kulis for critical reading of this manuscript.

## Author Contributions

X.A., F.P., S.B., D.C., E.C., contributed to sample collection as well as to their biological and clinical annotation; R.V-B., N.V-D., N.R., R.B., H.G.S., I.G., E.C., and J.I.M.-S. performed, coordinated and/or supported in situ Hi-C, histone mark, ATAC-seq, 4C-seq, methylome and transcriptome data generation; R.V-B., P.S-V., M.D-S., V.C., G.C., I.F., P.C., R.B., M.A.M-R., and J.I.M.-S. performed, coordinated and/or supported computational data analysis; R.V-B., P.S-V. M.D-S., I.F., P.C., E.C., M.A.M-R., J.I.M-S. participated in the study design and/or data interpretation. M.A.M-R. and J.I.M-S. directed the research and wrote the manuscript together with R.V-B. and P.S-V.

## Methods

### Isolation of B cell subpopulations for *in situ* Hi-C experiment

Four B cell subpopulations spanning mature normal B cell differentiation were sorted for *in situ* Hi-C as previously described^30^. Briefly, peripheral blood B cell subpopulations *i.e.* naive B cells (NBC) and memory B cells (MBC) were obtained from buffy coats for healthy adult male donors from 56 to 61 years of age, obtained from Banc de Sang i Teixits (Catalunya, Spain). Germinal center B cells (GCBC) and plasma cells (PC) were isolated from tonsils of male children undergoing tonsillectomy (from 2 to 12 years of age), obtained from the Clinica Universidad de Navarra (Pamplona, Spain). Samples were cross-linked before FACS sorting, to separate each of the B cell subpopulations, and afterwards were snap frozen and kept at −80°C. Three replicates per B cell subpopulation were processed and each replicate was derived from individual donors with the exception of plasma cells, for which two of the three replicates proceeded from the pool of four different donors. The use of the samples analyzed in the present study was approved by the ethics committee of the Hospital Clinic de Barcelona and Clinica Universidad de Navarra.

### Patient Samples

The samples from CLL (n=7)^28^ and MCL (n=5) patients were obtained from cryopreserved mononuclear cells from the Hematopathology collection registered at the Biobank (Hospital Clinic-IDIBAPS; R121004-094). All samples were >85% tumor content. Clinical and biological characteristics of the patients are shown in **Supplementary Table 5**.

The enrolled patients gave informed consent for scientific study following the ICGC guidelines and the ICGC Ethics and Policy committee^92^. This study was approved by the clinical research ethics committee of the Hospital Clinic of Barcelona.

### In situ Hi-C

*In situ* Hi-C was performed based on the previously described protocol^12^. Two million of cross-linked cells per sample were used as starting material. Chromatin was digested adding 100U Dpnll (New England BioLabs) on overnight incubation. After the fill-in with bio-dCTP (Life-Technologies, 19518-018), nuclei were centrifuged 5 minutes, 3000rpm at 4°C and ligation was performed for 4 hours at 16°C adding 2μl of 2000U/pl T4 DNA ligase on total 1.2mL of ligation mix (120μl of 10X T4 DNA ligase buffer; 100μl of 10% Triton X-100; 12μl of lOmg/ml BSA; 966μl of H_2_0). Following ligation, nuclei were pelleted and resuspended with 400μl 1X NEBuffer2 (New England BioLabs). Then, 10μl of RNAseA (lOmg/ml) was added to the nuclei and incubated during 15 minutes at 37°C while shaking (300rpm), and after that 20μl of proteinakse K (10mg/mL) was added and incubated overnight at 65°C while shaking (600rpm). After reversion of the cross-link, DNA was purified by phenol/chloroform/isoamyl alcohol and DNA was precipitated by adding to the upper aqueous phase: 0.1X of 3M sodium acetate pH 5.2, 2.5X of pure ethanol and 50μg/ml glycogen. Samples were mixed and incubated overnight at −80°C. Next, samples were centrifuged 30 minutes at 13,000rpm at 4°C and pellet was washed with lmL of EtOH 70% followed by a 15 minutes centrifugation at 13,000rpm at 4°C.

The supernatant was discarded and the pellet air-dried for 5 minutes and resuspended in 130μl of 1X Tris buffer (10 mM TrisHCI, pH 8.0), which to be fully dissolved was incubated at 37°C for 15 minutes. Purified DNA was sonicated using Covaris S220, and then the final volume was adjusted to 300μl with 1X Tris buffer. Sonicated DNA was mixed with washed magnetic streptavidin T1 beads (total of 100μl 10mg/ml beads), split in two tubes (150μl each), and incubated for 30 minutes at room temperature (RT) under rotation. Subsequently, beads were separated on the magnet, the supernatant discarded and the DNA was washed with 400μl of BB IX, twice. Sonicated DNA conjugated with beads was washed with 100μl of 1X T4 DNA ligase buffer, pooling the two tubes per condition. After that, beads were reclaimed in end-repair mix. Once incubated during 30 minutes at RT the beads were washed twice with 400μl of BB IX. Then, beads were washed with 100μl of NEBuffer2 and reclaimed in A-tailing mix, incubated during 30 minutes at 37°C and washed twice with 400μl of BB IX, followed by a wash in 100μl of 1X T4 DNA ligase buffer. Afterwards, the beads were resuspended in 50μl of 1X Quick ligation buffer, 2.5μl of lllumina adaptors and 4,000U of T4 DNA ligase and incubated during 15 minutes at RT. Then, beads were washed twice with 400μl BB 1X and resuspended in 30μl of 1X Tris buffer. In the end, libraries were amplified by eight cycle of PCR using 8.3μl of beads and pooling a total of 4 PCRs per sample. The PCR products were mixed by pipetting with an equal volume of AMPure XP beads and incubated at RT for 5 minutes. Beads were washed with 700μl of EtOH 70%, without mixing, twice, and left the EtOH evaporate at RT without over-drying the beads (aprox. 4 minutes). Finally, the beads were resuspended with 30μl 1X Tris buffer, incubated during 5 minutes and supernatant containing the purified library was transferred in a new tube and stored at −20°C. DNA was quantified by Qubit dsDNA High Sensitivity Assay, the library profile was evaluated on the Bioanalayzer 2100 and the ligation was assessed. Libraries were sequenced on HiSeq 2500. **Supplementary Table 1** summarizes the number of reads sequenced and quality metrics for each B cell subpopulation replicate and B cell neoplasm.

### Hi-C data pre-processing, normalization and interaction calling

The sequencing reads of Hi-C experiments were processed with TADbit^55^. Briefly, sequencing reads were aligned to the reference genome (GRCh38) applying a fragment-based strategy; dependent on GEM mapper^93^. The mapped reads were filtered to remove those resulting from unspecified ligations, errors or experimental artefacts. Specifically, we applied seven different filters using the default parameters in TADbit: self-circles, dangling ends, errors, extra dangling-ends, over-represented, duplicated and random breaks^55^. Hi-C data were normalized using the OneD correction^94^ at 100Kb of resolution to remove known experimental biases. The significant Hi-C interactions were called with the *analyzeHiC* function of the HOMER software suite^76^, binned at 10Kb of resolution and with the default p-value threshold of 0.001.

### Reproducibility of Hi-C replicas

The agreement between Hi-C replicates was assessed using the reproducibility score^36^. The RS is a measure of matrix similarity ranging between 0 (totally different matrices) and 1 (identical matrices). A genome-wide RS was defined for each experiment as the average RS between pairs of corresponding normalized chromosome matrix **(Extended Data Figure 1A** and **Extended Data 4B and 4D).** Then, the matrix representing all the genome-wide RSs was analyzed using a hierarchical clustering algorithm with the Ward’s agglomeration method using *hclust* function from R stats package.

### ChIP-seq and ATAC-seq data generation and processing

ChIP-seq of the six different histone marks and ATAC-seq data were generated as described in (http://www.blueprint-epigenome.eu/index.cfm?p=7BF8A4B6-F4FE-861A-2AD57A08D63D0B58)^28^. Briefly, fastq files of ChIP-seq data were aligned to the GRCh38 reference genome using bwa 0.7.7^95^, PICARD (http://broadinstitute.github.io/picard/) and SAMTOOLS^96^, and wiggle plots were generated (using *PhantomPeakQualTools* R package) as described (http://dcc.blueprint-epigenome.eu/#/md/methods). Peaks of the histone marks were called as described in http://dcc.blueprint-epigenome.eu/#/md/methods using MACS2 (version 2.0.10.20131216)^97^ with input control. ATAC-seq fastq files were aligned to genome build GRCh38 using bwa 0.7.7 (parameters: -q 5 -P -a 48 0)^95^ and SAMTOOLS vl.3.1 (default settings)^96^. BAM files were sorted and duplicates were masked using PICARD tools v2.8.1 with default settings (http://broadinstitute.github.io/picard/). Finally, low quality and duplicate reads were removed using SAMTOOLS vl.3.1 (parameters: -b -F 4 -q 5,-b, -F 1024)^96^. ATAC-seq peaks were determined using MACS2 (version 2.1.1.20160309, parameters: -g hs q 0.05 -f BAM -nomodel - shift -96 extsize 200 - keep -dup all) without input^97^.

For each mark a set of consensus peaks (chr1-22) present in the normal B cells (n=12 biologically independent samples for histone marks and n=15 biologically independent samples for ATAC-seq) was generated by merging the locations of the separate peaks per individual sample. Also, a second set of consensus peaks was generated taking into account normal B cells, CLL (n=7 biologically independent samples) and MCL (n=5 biologically independent samples). For the histone marks, the number of reads per sample per consensus peak was calculated using the *genomecov* function of bedtools suite^98^. For ATAC-seq, the number of insertions of the TN5 transposase per sample per consensus peaks was calculated determining the estimated insertion sites (shifting the start of the first mate 4bp downstream), followed by the *genomecov* function of bedtools suite^98^. The number of consensus peaks for normal B cell samples were 46,184 (H3K4me3), 44,201 (H3K4me1), 72,222 (H3K27ac), 25,945 (H3K36me3), 40,704 (H3K9me3), 20,994 (H3K27me3), 99,327 (ATAC-seq), while the number of consensus peaks for normal B cells, CLL and MCL samples were 53,241 (H3K4me3), 54,653 (H3K4me1), 106,457 (H3K27ac), 50,530 (H3K36me3), 137,933 (H3K9me3), 117,560 (H3K27me3), 140,187 (ATAC-seq). Using DESeq2 R package^99^, counts for all consensus peaks were transformed by means of the variance stabilizing transformation (VST) with blind dispersion estimation. Principal component analysis (PCAs) were generated with the *prcomp* function from the stats package in R using the VST values.

### RNA-seq data generation and processing

Single-stranded RNA-seq data were generated as previously described^100^. Briefly, RNA was extracted using TRIZOL (Life Technologies) and libraries were prepared using TruSeq Stranded Total RNA kit with Ribo-Zero Gold (Illumina). Adapter-ligated libraries were amplified and sequenced using 100bp single-end reads. RNA-seq data of the 24 samples, some (n=19) mined from a previous study^28^, were aligned to the reference human genome build GRCh38 (**Supplementary Table 5**). Signal files were produced and gene quantifications (gencode 22, 60,483 genes) were calculated as described (http://dcc.blueprint-epigenome.eu/#/md/methods) using the GRAPE2 pipeline with STAR-RSEM profile (adapted from the ENCODE Long RNA-Seq pipeline). The expected counts and fragments per kilobase million (FPKM) estimates were used for downstream analysis. The PCA of the RNA-seq data was generated with the *prcomp* function from the stats package in R in the 12 analyzed normal B cell samples or 24 analyzed normal and neoplastic B cell samples.

### WGBS data generation and processing

WGBS was generated as previously described^30^. Mapping and determination of methylation estimates were performed as described (http://dcc.blueprint-epigenome.eu/#/md/methods) using GEM3.0. Per sample, only methylation estimates of CpGs with ten or more reads were used for downstream analysis. The principal component analysis (PCA) of the DNA methylation data was generated with the *prcomp* function from the stats package in R using methylation estimates of 15,089,887 CpGs (chr1-22) with available methylation estimates in all 12 analyzed normal B cell samples or 14,088,025 CpGs (chr1-22) in all 24 analyzed normal and neoplastic B cell samples.

### Definition of sub-nuclear genome compartmentalization

The segmentation of the genome into compartments was determined as previously described^5^. In short, normalized chromosome-wide interaction matrices at 100Kb resolution were transformed into Pearson correlation matrices. These correlation matrices were then used to perform PCA for which the first eigenvector (EV) normally delineates genome segregation. All EVs were visually inspected to ensure that the EV selected corresponded to genomic compartments^5^. Since the sign of the EV is arbitrary, a rotation factor based on the histone mark H3K4me1 signal and ATAC-seq signal were applied to correctly call the identity of the compartments. A Pearson correlation coefficient was computed between the EVs for each pair of merged B cell subpopulation **(Extended Data Figure 1C).** Each merged sample was also correlated with its replica **(Extended Data Figure 1C).** The multi-modal distribution of the EV coefficients from the B cells dataset was modelled as a Gaussian mixture with three components (k = 3). To estimate the mixture distribution parameters, an Expectation Maximization algorithm using the *normalmixEM* function from the *mixtools* R package was applied^101^.

A Bayesian Information Criterion (BIC) was computed for the specified mixture models of clusters (from 1 to 10) using *mclustBIC* function from *mclust* package in R^102^ **(Extended Data Figure 1D).** Three underlying structures were defined; an alternative compartmentalization into A-type (with the most positive EV values), B-type (with the most negative EV values) and I-type (an intermediate-valued region with a distinct distribution) compartments. Two intersection values (IV1, IV2) were defined at the intersection points between two components. The mean IV1 and IV2 values across all the B cell replicas (n=12) were then used as standard thresholds to categorize the data into the three different components (that is, A-type compartment was defined for EV values between +1.00 and +0.63, I-type compartment as of “Intermediate” was defined for EV values between +0.63 and −0.43, and B-type compartment was defined for EV values between −0.43 and −1.00) **(Extended Data Figure 1E).**

### Characterizing compartment types in B cells by integrating nine omics layers

Given a set of peaks as previous defined by Beekman et al^28^ from nine different omic layers including six histone marks (H3K4me3, H3K4me1, H3K27ac, H3K36me3, H3K9me3, H3K27me3), gene accessibility (ATAC-seq), gene expression (RNA-seq) and DNA methylation (WGBS), a *bedmap* function from BEDOPS software^103^ was applied to get the mean scoring peak over the 100Kb intervals genome-wide. Next, Pearson correlation coefficients were computed between the EV coefficients and the mean scoring value of each epigenetic mark at 100Kb intervals **(Extended Data Figure 1B).** Finally, the mean scoring values were normalized by the total sum of the values for each mark and grouped by the three defined genomic compartments (A, I, B-type; Figure 1G). A Wilcoxon test was used to compute the significance between all the possible pairwise comparisons of the signal distribution.

### Compartment Interaction Score (C-Score)

The compartment score is defined as the ratio of contacts between regions within the same compartment (intra-compartment contacts) over the total chromosomal contacts per compartment (intra-compartment + inter-compartment). To compute the compartment score, all the compartments that shared the same genomic segmentation were merged.

### Chromatin states enrichment by genomic compartments

The genome was segmented into 12 different chromatin states at 200bp interval as previously described^28^. The active promoter and strong enhancerl were merged as a unique state, giving a total of 11 chromatin states. The genome compartmentalization was next split into 4 groups; 3 conserved groups, in which the B cells samples shared A-type compartment (n=6,409), B-type compartment (n=6,267) or I-type compartment (n=5,467) and a dynamic group (n=7,099) of non-conserved compartmentalization among B cells subpopulation. Each group was correlated with the defined 11 chromatin states using *foverlaps* function from *data.table* R package. The frequency of each chromatin state (corrected by the total frequency in the genome) was computed per each genomic compartment. The chromatin state score is thus the median frequency of the three replicas scaled by the columns and the rows using *scale* function from *baseR* package.

### Description of chromatin states in the intermediate (I)-type compartment

200bp-windows containing poised promoter (n=547) or polycomb repressed (n=11,665) chromatin states were extracted from the NBC intermediate compartments (n=l,885). From those regions, two main sub-groups were distinguished according to the chromatin state shown in the next state of differentiation (GCBC): (1) those regions that maintained their chromatin state (poised promoter or polycomb repressed), and (2) those regions that changed their chromatin state; which were further classified into three categories: (i) l-related chromatin states (poised promoter or polycomb repressed), (ii) B-related chromatin states (repressive heterochromatin and low signal heterochromatin), (iii) A-related chromatin states (active promoter/strong enhancer1, weak promoter, strong enhancer2, transcription transition, transcription elongation and weak transcription). Finally, the fold-change of related chromatin states between GCBC and NBC was computed.

### Analysis of chromatin state dynamics upon B cell differentiation

B cell differentiation axis was divided into two main branches: (i) NBC-GCBC-PC, (ii) NBC-GCBC-MBC. Both branches presented a common step from NBC to GCBC and then a divergence step in PC or MBC. The 5,445 common compartments from both branches were considered for the analysis. The general modulation of chromatin structure was drawn using the *alluvial* function from *alluvial* R package.

### Transcription factor analyses

From GCBC-specific 937 active compartments (B to A-type, n =18; B to I-type, n=512 and I to A-type, n=407) were narrowed down to 171 peaks due to the following filtering steps: (i) only the 200bp-windows contain active promoter, strong enhancerl and strong enhanced chromatin states were retained (n=1,907 regions), (ii) Regions where H3K27ac peaks were differentially enriched in GCBC replicates compared to the rest of normal B cell subpopulations (FDR<0.05) computed using DEseq2 R package” were retained, (iii) Regions with a presence of ATAC-seq peaks in at least two GCBC replicates were retained (n=171 peaks). The background considered was the rest of the ATAC-seq peaks (n=268) presented at the 1,907 regions in at least two GCBC replicates.

From CLL-specific 48 active compartments (in normal B cells defined as I-type: n=28 and B-type: n=20), were narrowed down to 25 peaks due to the following filtering steps: (i) Regions where H3K27ac peaks were differentially enriched (FDR<0.05) comparing CLL from all normal B cells and MCL using DEseq2 package”, (ii) Regions where ATAC-seq peaks were presented in at least five CLL (n=25). The background considered was all the resting ATAC-seq peaks (n=28) on the 48 compartments presented in at least five CLL.

On both analysis, FASTA sequences of targeted regions (GCBC-specific regions and CLL-specific regions) were extracted using *getfasta* function from *bedtools* suite^98^ using GRCh38 as reference assembly. An analysis of motif enrichment was done by the *AME-MEME* suite^104^ using non-redundant transcription factor (TF) binding profiles of *Homo sapiens* Jaspar 2018 database^105^ as a reference motif database. The database contained a set of 537 DNA motifs. Maximum odd scores were used as a scoring method and one-tailed Wilcoxon rank-sum as motif enrichment test. Only TF genes that were expressed (FPKM median values>1) were included.

### TCF4 binding motif example from *KSR2* gene

A FASTA sequences of 25 ATAC-seq peaks detected in CLL-specific active compartments were extracted using GRCh38 as reference assembly. A search of individual motif occurrences analysis was done using *AME-FIMO* suite^106^ library(BSgenome.Hsapiens.UCSC.hg38,masked) with a custom random model (letter frequencies: *A,* 0.262: *C,* 0.238: G, 0.238 and T, 0.262). A p-value<0.0001 was established as a threshold to determine 23 significant motif occurrences where TCF4 binding motif (MA0830.1) was one of the top candidates.

### Log-ratio of normalized interactions in the *AICDA* regulatory landscape

Normalized Hi-C maps were analyzed at 50Kb of resolution at the specific genomic region, chr12:8,550,000-9,050,000 (GRCh38), from the four B cell subpopulations. A logarithmic ratio of the contact maps was computed between NBC and GCBC and GCBC with PC and MBC. The result array was convolved with a 1-dimensional Gaussian filter of standard deviation (sigma) of 1.0 using and interpolated with a nearest-neighbor approach using *scipyndimage* Python package.

### Statistical testing for detecting significant changed compartment regions

Briefly, 100Kb regions that had at least one missing value among the compared samples were removed from the analysis. Then, two different groups were defined, case and control, according to the case-control pair analyzed. A T-test was computed to compare each case-control pair, and the resulting p-values were adjusted using the false discovery rate (FDR)^107^. The regions with significantly different means and fold changes were selected based on two specific thresholds: a p-adjustment value less than 0.05 and a fold change greater than 0.4. The results were then generated for a total of 4 different case-control pairs.

(I) control: all regions conserved across all B cell samples without missing values in CLL (A-type, n=3,967, I-type, n=4,301 and B-type, n=5,226), case: all CLL regions non-conserved in B cell samples (n=3,217). The analysis resulted in 348 B cell_CLL significantly changed regions.

(II) control: all regions conserved across all B cell samples without missing values in MCL (A-type n=6,167, I-type n=5,299, B-type n=5,812), case: all MCL regions non-conserved in B cell samples (n=4,716). The analysis resulted in 82 B cell_MCL significantly changed regions.

(III) control: B cell-CLL significantly changed regions (n=348) - MCL-CLL overlapping (n=31) = B cell-CLL specific regions (n=317), case: MCL regions (A-type n=97, I-type n=154, B-type n=61; total n=312). The analysis resulted in 89 B cell_CLL-specific regions.

(IV) control: B cell-MCL significantly changed regions (n=82) - MCL-CLL overlapping (n=31) = B cell-MCL specific regions (n=51), case: CLL regions (n=41). The analysis resulted in 3 B cell_MCL-specific regions.

### Integrative 3D modelling of EBF1 and structural analysis

Hi-C interactions matrices from the merging of three replicas of NBC and the seven cases of CLL were used to model chr5:158,000,000:160,000,000 (GRCh38) at 5Kb of resolution. For NBC and CLL merged Hi-C interaction maps, a MMP score was calculated to assess the modeling potential of the region, resulting in 0.79 for NBC and 0.84 for CLL indicative of good quality Hi-C contact maps for accurate 3D reconstruction^108^. Next, this region was modelled using a restraint-based modelling approach as implemented in TADbit^55^, where the experimental frequencies of interaction are transformed into a set of spatial restraints^54^. Briefly, each 5Kb bin of the interaction Hi-C map was represented as a spherical particle in the model, which resulted in 400 particles each of radius equal to 25nm. All the particles in the models were restrained in the space based on the frequency of the Hi-C contacts, the chain connectivity and the excluded volume. The TADbit optimal parameters (maxdist=-1.0; lowfreq=1.0; upfreq=200; and dcutoff=150) resulted in the best Spearman correlations of 0.61 (NBC) and 0.63 (CLL) between the Hi-C interaction map and the models contact map. Next, a total of 5,000 models per cell type were generated, and the top 1,000 models that best satisfied the imposed restraints were retained for the analysis. To assess the structural similarities among the 3D models, the distance root-mean-square deviations (dRMSD) value was computed for all the possible pairs of top models (1,000 in NBC and 1,000 in CLL) and a hierarchical clustering algorithm was applied on the resulting dRMSD matrix using *ward.D* method from stats package in R **(Extended Data Figure 5C).** The convex hull volume spanned by the 81 particles of the *EBF1* gene (chr5:158,695,000-159,000,000, GRCh38) was computed in each model using the *convexhull* function from the *scipy.spatial* Python package (Figure 5G).

### Differential Gene expression analyses

Differentially expressed genes were defined using the DEseq2 R package”, *nbinomWaldTest,* on all the genes. Then, the genes present on the compartments of interest were selected and Benjamini y Hochberg (BH) test (FDR<0.05) was applied. In detail, expected counts were used on the following considered comparisons: (i) for GCBC-specific activate compartments, GCBC samples (n=3) versus the rest of normal B cells samples (NBC, PC, MBC; n=9); (ii) for CLL-specific active compartments, CLL samples (n=7) versus the rest of the samples (normal B cells and MCL, n=17); (iii) for CLL-specific inactive compartments, all normal B cells and MCL samples (total n=17) versus CLL samples (n=7) and (iv) for cMCL, cMCL (n=2) versus nnMCL (n=3) samples were studied. Then, the expression of the genes differentially expressed on each comparison of interest was assessed. Only genes that were expressed (FPKM median values>l) were included.

The *findOverlaps* function from *GenomicRanges* R package^109^ was used to annotated genes that overlapped with these defined regions. One tailed Monte-Carlo method was applied to evaluate the significant number of differentially expressed genes in CLL-specific compartments (this process was randomly repeated 10,000 times).

### Defining de novo (in)active regions in sub-type specific neoplastic group

MCL and CLL patient samples were grouped according to their biological and clinical characteristics. This classification resulted in two conventional (c) and three leukemic non-nodal (nn) MCL cases and two IGVH-unmutated (u) and five IGVH-mutated (m) CLL cases.

First, the non-assigned neoplasia compartments were removed from the analysis. A sample homogenization was applied to reduce the intra-subtype variance; the samples that presented a difference of EV smaller than 0.4 were retained (91.29% in MCL, 87.1% CLL). Next, to study the inter-subtype variance, the mean of the EV from each subtype of B cell malignancy was computed. Significant regions were determined if the difference between the two subtypes (cMCL vs nnMCL and uCLL vs mCLL) was equal or higher than 0.4, which resulted in 673 regions in MCL and 47 in CLL. MCL-subtype specific regions where split into two groups according to the value of its EV coefficient (n=435 region called cMCL gain, n=238 regions called nnMCL gain). The distribution and the frequency of the significantly changed regions were studied per chromosome and compared with the probability of finding them by chance in each chromosome. N-subsamples of 100Kb size were selected from the GRCh38 genome and their frequency was calculated per chromosome (this process was randomly repeated 10,000 times). One tailed Monte-Carlo method was applied to compute p-values. The *findOverlaps* function from *GenomicRanges* R package (Lawrence et al. 2013) was next used to annotate protein coding genes that overlapped with these defined regions. Differentially expressed genes among cMCL and nnMCL on chr2:2,700,000-8,800,000 (GRCh38) was compute using *Deseq2* (Love et al. 2014) (using a FDR<0.05). The expression analysis was validated on two independent published cohorts, i.e.: a series with 30 conventional and 24 leukemic non-nodal mantle cell lymphoma (GEO GSE79196) from peripheral blood^49^ and a second series from the lymphoma/leukemia molecular profiling project (LLMPP) (GEO GSE93291)^62^. The microarrays were normalized using the R *frma* method and *Umma* R package was used to identify differentially expressed genes with adjusted p-value<0.05. Standardized expression matrices were used to do the heatmaps using *pheatmap* R package. Gene differentially expressed on the identified cohort: [1] RNAseq from BLUEPRINT data, [2] peripheral blood and [3] LLMPP. The magnitude of the compartmentalization change was calculated subtracting the EV of cMCLl and nnMCL2. The karyotype and the chromosome 2 were designed using the *karyoploteR* library^112^.

**Extended Data Figure 1**

**Extended Data Figure 1A.** Average genome-wide reproducibility score matrix of each B cell subpopulation replicate at 100Kb resolution. The reproducibility score ranging between 0 (totally different matrices) and 1 (identical matrices).

**Extended Data Figure 1B.** Pearson correlation between the eigenvector coefficients, which defines 3D compartments per B cell subpopulation, with six histone marks, chromatin accessibility (ATAC-seq), gene expression (RNA-seq) and DNA methylation (WGBS). Positive values of the eigenvector show higher correlation with H3K4me1 (enhancer mark) and chromatin accessibility.

**Extended Data Figure 1C.** Genome-wide scatterplots of coefficients from the first eigenvector showing the correlation between pairs of B cell subpopulations at 100Kb resolution. The squared correlation coefficient (R^2^) is indicated.

**Extended Data Figure 1D.** Bayesian Information Criterion (BIC) plot for the equal (E) and unequal (V) variance model parameterization ranged from 1 to 10 clusters.

**Extended Data Figure 1E.** Compartment definition model. The x-axis shows the distribution of the eigenvector coefficients and the y-axis indicates the density. The fitting model proposed is highlighted using solid black line. The red lines mark the intersection points (EV1 = −0.63 and EV2 = 0.43) used to distinguish the three different compartments (A-type, I-type, B-type).

**Extended Data Figure 2.**

**Extended Data Figure 2A.** Functional validation of the conserved (A-type, I-type and B-type) and dynamic compartments in all B cell subpopulations replicates using eleven different chromatin states. The chromatin state score is normalized by sample and chromatin state.

**Extended Data Figure 2B.** C-score. Method defined by the ratio of contacts betwwen regios within the same compartment (intra-compartment contacts) over the total chromosomal contacts per compartments (intra- and inter-chromosomal interactions).

**Extended Data Figure 2C.** C-score distributions on the three defined compartments A-type, I-type and B-type.

**Extended Data Figure 2D.** C-score distributions segmenting the I-type compartment onto positive (IA) or negative (IB) eigenvector coefficients.

**Extended Data Figure 4.**

**Extended Data Figure 4A.** Average genome-wide reproducibility score matrix of each B cell subpopulation replicate and B cell neoplasia at 100Kb. The reproducibility score ranging between 0 (totally different matrices) and 1 (identical matrices).

**Extended Data Figure 4B/4D.** Genome-wide scatterplots of the first eigenvector showing the correlation between pairs of each B cell malignancy samples at 100Kb resolution. CLL **(B).** MCL **(D).** The squared correlation coefficient (R^2^) is indicated.

**Extended Data Figure 4C/4E.** Mean and standard deviation of the squared correlation coefficients calculated intra- or inter-each neoplasia subtype. CLL **(C).** MCL **(E).**

**Extended Data Figure 5.**

**Extended Data Figure 5A.** Heatmap showing the eigenvector coefficients of the compartments losing **(top)** or gaining **(bottom)** activation specifically in MCL.

**Extended Data Figure 5B.** Correlation between normalized Hi-C and modeled contact maps in *EBF1* regulatory landscape. **Left:** Contact map computed from the restrained-based model. **Middle:** Scatterplot of Hi-C normalized map versus modeled contact data with linear regression. **Right:** Normalized Hi-C data. **Top:** NBC. **Bottom:** CLL. The position of *EBF1* is indicated in blue at the bottom of the matrix plots.

**Extended Data Figure 5C.** Heatmap of the hierarchical clustering of the dRMSD values computed for all the possible pairs of generated models (1,000 in NBC and 1,000 in CLL).

**Extended Data Figure 7.**

**Extended Data Figure 7A.** Bar graphs represent the fold change between cMCL and nnMCL of each three groups of chromatin states (arranged by their relationship to the A-type, I-type and B-type compartments). Active Promoter, Weak Promoter, Strong Enhancer 1, Strong Enhancer 2, Weak Enhancer, Transcription Transition, Transcription Elongation, Weak Transcription were A-type compartment-related states. Heterochromatin;Repressed and Heterochromatin;Low signal were B-type compartment-related states. Poised Promoter or Polycomb repressed chromatin states were I-type compartment-related states.

**Extended Data Figure 7B/7C.** Heatmaps of the differentially expressed gens between MCL samples classified as cMCL (light yellow) and nnMCL (dark yellow) subtypes. Peripheral blood **(B)** and LLMPP **(C)** cohorts. The VST values were normalized by genes.

**Supplementary Tables**

**Supplementary Table 1.** *In situ* Hi-C experimental quality metrics.

**Supplementary Table 2.** GCBC specific 3D active compartments on a three-column bed file format (chromosome, start position and end position).

**Supplementary Table 3.** List of the identified enriched binding motifs expressed on GCBC.

**Supplementary Table 4.** Genes differentially upregulated (FDR<0.05) in GCBC specific regions. The coordinates of the compartment or compartments each gene belongs to is indicated

**Supplementary Table 5.** Patient characteristics and general overview of the omic layers analyzed.

**Supplementary Table 6.** Genes differentially expressed (FDR<0.05) at CLL-specific inactive compartments. The coordinates of the compartment or compartments each gene belongs to is indicated

**Supplementary Table 7.** Genes differentially expressed (FDR<0.05) at CLL-specific active compartments. The coordinates of the compartment or compartments each gene belongs to is indicated

**Supplementary Table 8.** List of the identified enriched binding motifs expressed on CLL-specific active compartments.

