## Supplementary figures and images for "Dynamics of genome architecture and chromatin function during human B cell differentiation and neoplastic transformation" (from another version of this preprint: version 1, 2019-09-10)

### Extended Figure 5

#### Extended Data Figure 5

A

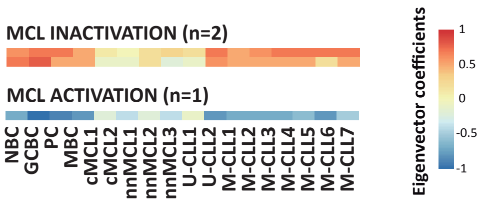


B

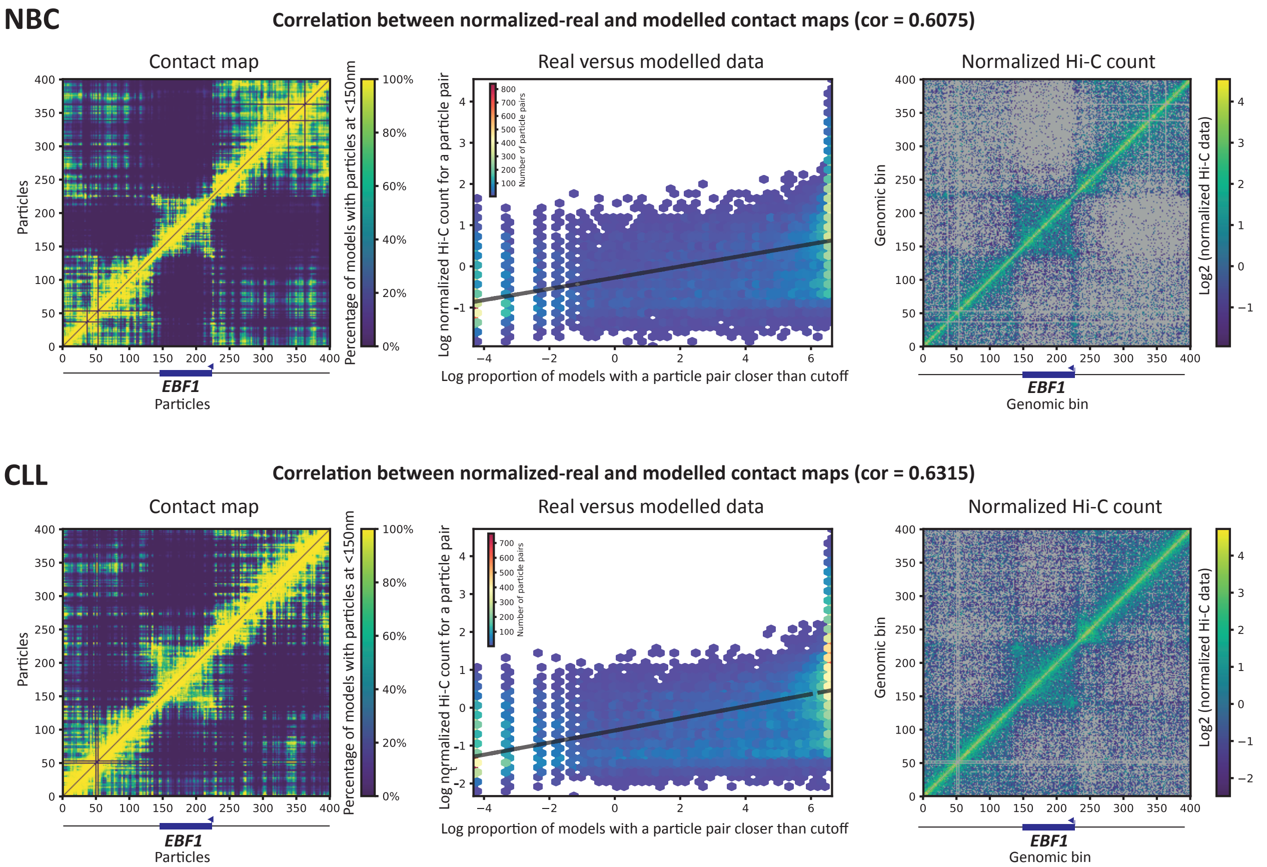


C

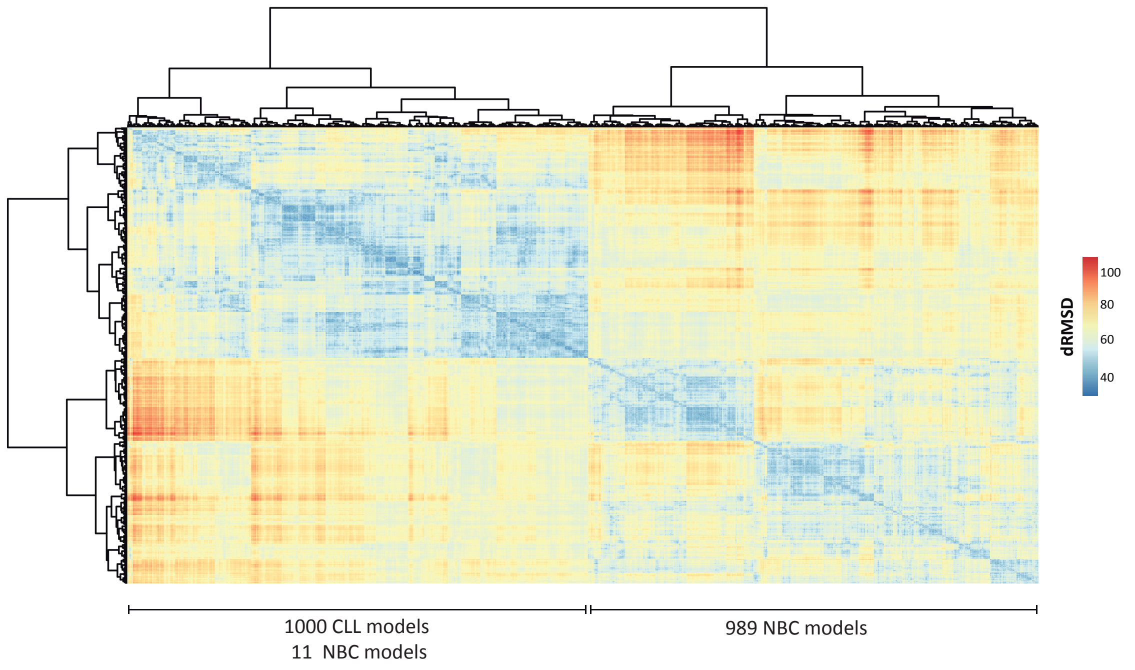
